# Sequential division of labor between PAXX and XLF drives NHEJ synaptic complex stability and maturation

**DOI:** 10.64898/2026.08.10.743781

**Authors:** Jinghang Zhang, Philippe Frit, Andrew T. Moreno, Nadia Barboule, Patrick Calsou, Sébastien Britton, Joseph J. Loparo

**Affiliations:** Department of Biological Chemistry and Molecular Pharmacology, Blavatnik Institute, Harvard Medical School, Boston, MA 02115, USA; IPBS, CNRS, Univ Toulouse, Toulouse, 31077, France

## Abstract

DNA double-strand breaks (DSBs) are primarily repaired by non-homologous end joining (NHEJ), which tethers and ligates DNA ends within a synaptic complex. PAXX is an NHEJ accessory factor related to XRCC4 and XLF, but its mechanistic role and genetic interaction with XLF are unclear. Using biochemical assays and single-molecule imaging within *Xenopus laevis* egg extracts and human cell assays, we show that PAXX bridges DNA ends by using its disordered tails to bind opposing Ku molecules. This PAXX tether substantially extends long-range synaptic complex (LRSC) lifetime and modestly stabilizes the short-range synaptic complex (SRSC). LRSC stabilization increases XLF residence and promotes the LRSC-to-SRSC transition. Using domain-swapped chimeras, we demonstrate that PAXX’s tails provide synaptic stabilization, whereas XLF’s heads interact with XRCC4 to drive the LRSC-to-SRSC transition: an XLF head-PAXX tail chimera recapitulates the functions of both proteins, revealing complementary, sequential roles for PAXX and XLF in NHEJ.

## Introduction

DNA double-strand breaks (DSBs) are among the most deleterious forms of DNA damage. If left unrepaired, even a single DSB can cause cell death, while misrepaired DSBs can produce oncogenic mutations and chromosomal translocations^1^. In addition to spontaneous DSBs that result from exogenous and endogenous agents, vertebrates must repair programmed DSBs generated during adaptive immune diversification^2,3^. To combat the detrimental effects of unrepaired DSBs, cells have evolved multiple DSB repair pathways. In humans, homologous recombination (HR) and non-homologous end joining (NHEJ) are the two major pathways. HR uses a sister chromatid as a homologous template to perform largely error-free repair and is therefore restricted to S and G2 phases. In contrast, NHEJ is the dominant DSB repair pathway because it directly ligates broken DNA ends together and occurs throughout the cell cycle^1^.

NHEJ is executed by a set of core factors consisting of the Ku70/Ku80 heterodimer (Ku)^4^, DNA-dependent protein kinase catalytic subunit (DNA-PKcs)^5–8^, X-ray cross complementing factor 4 (XRCC4)^8–11^, DNA ligase IV (Lig4)^8,10,12,13^ and XRCC4-like factor (XLF)^8,10,11,14^. Biochemical reconstitutions of these factors established that they are sufficient to assemble a synaptic complex which bridges the DNA ends and ligates them^6,15–17^. Complex DNA termini require additional processing by NHEJ-associated polymerases, nucleases and other enzymes before ligation^1,18,19^. In addition, accessory factors, including Paralog of XRCC4 and XLF (PAXX)^17,20–23^, MRI^24^, APLF^25–27^, and WRN^28–30^, stimulate NHEJ, most likely by promoting DNA end synapsis through still ill-defined mechanisms.

Formation and maintenance of a stable DNA end synaptic complex, which evolves during repair, is critical for accurate NHEJ^6,18,23,31,32^. First, DNA ends pair in a long-range synaptic complex (LRSC), in which the ends are held ∼100 Angstroms apart. LRSC formation requires Ku and DNA-PKcs, with DNA-PKcs providing the primary synaptic interface^8,31^. Multiple LRSCs^23,32^ have been visualized by structural studies, although their functional roles remain unclear. Transition to a short-range synaptic complex (SRSC) aligns the DNA ends for ligation and requires XRCC4, XLF, and Lig4^8,31^. Because long-range synapsis is transient, mechanisms that extend LRSC lifetime may be critical to provide a kinetic window for assembly of factors that drive synaptic maturation to the long-lived and ligation-competent SRSC. DNA end processing is largely confined to the SRSC, which prioritizes ligation of compatible ends and minimizes sequence alteration^18^. Therefore, stable synapsis is key to ensure high-fidelity NHEJ, as loss of synapsis not only precludes downstream end processing and ligation, but subsequent mispairing of ends results in potentially oncogenic translocations^1^.

PAXX is a recently identified NHEJ accessory factor that is structurally related to XRCC4 and XLF^20–22^. Like its homologs, PAXX forms a homodimer with each monomer comprising an N-terminal globular domain, a helical dimerization domain, and a long, disordered C-terminal tail. The C-termini of the tails of PAXX and XLF bear Ku binding motifs (KBMs) that interact with Ku70 and Ku80, respectively^20,21,23,33,34^ (Figure 1A). Both XRCC4 and XLF have well-defined roles in NHEJ. XRCC4 forms a stable complex with Lig4 that is essential for Lig4 stability^35^, and it interacts with XLF via a head-to-head interaction that is required for SRSC formation^8,10,14,31^. Consistent with these central roles, XRCC4 loss causes profound DSB repair defects and embryonic lethality^36,37^. By contrast, loss of either PAXX or XLF results in less severe phenotypes. Though XLF loss in humans results in clinical features such as immunodeficiency and microcephaly, human embryos deficient in XLF are still viable^38^, and XLF-knockout mice show extremely mild phenotypes at most ^39–42^. PAXX-deficient mice also show at most mild phenotypes ^39–41^, and there is no known phenotype associated with PAXX deficiency in humans. However, combined loss of both factors in mice causes embryonic lethality, suggesting substantial functional redundancy^39–41^. Together with the established role of XLF in DNA end synapsis, these observations suggest that PAXX also contributes to synapsis^6^. However, they do not define the stages of NHEJ at which PAXX acts, how its structural features support DNA end tethering, or how its function overlaps with or complements that of XLF.

**Figure 1:**
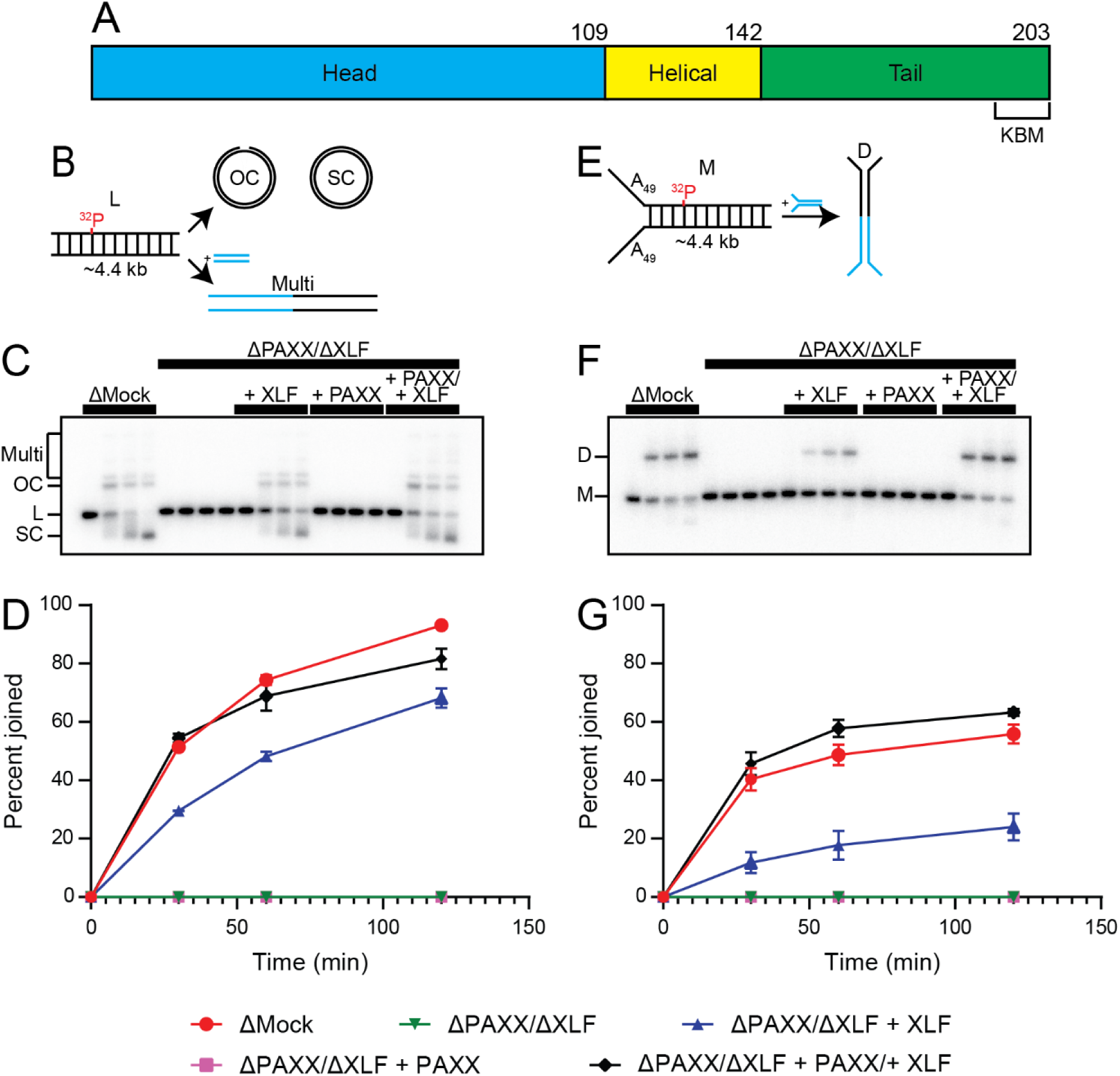
PAXX stimulates NHEJ in *X. laevis* egg extract. (A) Domain architecture of wild type PAXX (monomer). (B) Schematic depicting substrate and products for the double blunt ended substrate. L: linear substrate, OC: open circular singly-ligated intermediate, SC: supercoiled doubly-ligated product, Multi: multimeric product. (C) Agarose gel showing products of end joining reaction using substrate in Figure 1B. (D) Quantification of Figure 1C. (E) Schematic depicting substrate and products for the forked end substrate. M: monomeric substrate, D: dimeric product. (F) Agarose gel showing products of end joining reaction using substrate in Figure 1E. (G) Quantification of Figure 1F. Error bars in Figure 1D and Figure 1G represent ± SEM.

Here, we combine biochemical and single-molecule experiments in *Xenopus laevis* egg extract and human cell-based assays to define how PAXX promotes NHEJ. We first show that PAXX depletion from egg extract impairs, but does not abolish, end joining. Single-molecule fluorescence colocalization and FRET analysis reveal that PAXX primarily extends LRSC lifetime, modestly stabilizes the SRSC, and increases XLF residence at DNA ends, which promotes the LRSC-to-SRSC transition. Motivated by structural studies showing that the PAXX C-terminal tails can simultaneously engage opposing Ku molecules^23^, we tested whether shortening these tails or disrupting one or both KBMs compromises PAXX function. We reveal that these perturbations abrogate PAXX-mediated stimulation of NHEJ, supporting a model in which PAXX’s tails tether to Ku dimers on opposite DNA ends to bridge and stabilize the synaptic complex. To define the shared and distinct functions of PAXX and XLF in NHEJ, we constructed chimeras of both proteins in which we exchanged their tail domains. A chimera comprising the XLF head/helical domains fused to PAXX tails successfully rescued a combined PAXX/XLF depletion, whereas the reciprocal chimera did not, indicating that PAXX’s principal function is to stabilize synapsis via its tails, with its ordered domains serving primarily to dimerize these modules. Conversely, XLF promotes LRSC-to-SRSC transition through head–XRCC4 interactions, while its tail domains primarily mediate recruitment to the synaptic complex. Together, these results establish how PAXX-dependent end synapsis promotes XLF incorporation and SRSC formation, explain the mechanistic basis for the observed redundancy between XLF and PAXX within NHEJ, and suggest a general mechanistic framework for the several pairwise redundancies observed between various other NHEJ factors.

## Results

### PAXX is required for efficient NHEJ in Xenopus egg extract

To determine the importance of PAXX in NHEJ, we used *Xenopus laevis* egg extract, which carries out end joining in a manner that requires the core NHEJ factors^43–46^. We performed end joining assays by incubating a ∼4.4 kb blunt-ended ^32^P-radiolabeled DNA substrate in egg extract. End joining products were separated at distinct time points by running the recovered DNA through an agarose gel. Over the course of the reaction, the linear substrate primarily underwent intramolecular end joining in which open circular intermediates were ultimately converted to supercoiled products (Figure 1B, C). Multimers of the input substrate were also observed as a minor product (Figure 1C). Egg extract was immunodepleted of PAXX using an antibody raised against full length *X. laevis* PAXX (Extended Data 1C). Given that our anti-PAXX antibody also partially depleted XLF from the extract (Extended Data 1C), we co-depleted XLF using an anti-XLF antibody and supplemented the extract with recombinant *X. laevis* XLF to control the exact amount of XLF in our reactions. Unless otherwise indicated, all further uses of “PAXX depletion” or “PAXX depleted extract” in this work refer to PAXX/XLF double depleted extract supplemented with 50 nM recombinant XLF to simulate a single depletion of PAXX. This concentration of XLF was sufficient to rescue XLF depleted extracts to levels of end joining comparable to extract that underwent a mock depletion using non-specific rabbit IgG (Extended Data 1D, E). Consistent with our previous results demonstrating that XLF is essential for NHEJ in egg extract, depletion of both PAXX and XLF, with or without rescue with recombinant PAXX, abolished end joining activity. Rescue with only recombinant XLF partially restored NHEJ. However, NHEJ in the absence of PAXX was still modestly defective compared to NHEJ in mock depleted extract, and a full rescue only occurred with the addition of physiological concentrations of both PAXX and XLF (Figure 1C, D). These results demonstrate that PAXX is required for robust and efficient NHEJ in egg extract.

Because PAXX has been implicated in DNA end synapsis during NHEJ^6^, we hypothesized that intermolecular end joining may be more dependent on PAXX than intramolecular end joining. To that end, we designed a fork-containing DNA substrate that can only form a dimeric product mediated by a single intermolecular end joining event. This substrate is identical to the intramolecular joining substrate described above except for the addition of a fork comprising two 49 bp poly-A ssDNA tails at one end, which prevents Ku loading and ensures that each substrate molecule undergoes only a single end joining event at the blunt end (Figure 1E, F). As with the double-blunt ended substrate, NHEJ was abolished in the absence of XLF and defective when only PAXX was absent, with full rescue occurring only when XLF and PAXX were both present (Figure 1F, G). Consistent with our hypothesis, the relative end joining defect at the end of the time course in the absence of PAXX was much greater for the forked end substrate (>50% defect in PAXX depleted extract relative to mock depleted extract) than for the blunt ended substrate (∼25% defect in PAXX depleted extract relative to mock depleted extract) (Figure 1C-D, F-G). This difference in end joining suggests that PAXX promotes the early stages of DNA end synapsis, as its presence is less essential for an intramolecular substrate where ends are present at a high local concentration. Collectively, these ensemble end-joining assays show that PAXX enhances NHEJ efficiency, and that its contribution is particularly important for substrates constrained to intermolecular joining, consistent with a primary role in promoting early DNA end synapsis.

### PAXX stabilizes both the long-range and short-range synaptic complexes and facilitates synaptic transition

To determine how PAXX contributes to DNA end synapsis, we used single-molecule fluorescence assays developed by our laboratory^8^ to observe the formation and dissociation of the LRSC and SRSC. We first measured the stability of the LRSC using an intermolecular synapsis assay. For this assay, we attached a blunt-ended DNA duplex, which was biotinylated at one end and labeled with a donor fluorophore near the other end, to a streptavidin coated coverslip within a microfluidic flow cell (Figure 2A). To observe synapsis with the tethered DNA molecules, we then added an acceptor fluorophore-labeled DNA duplex in solution in which one DNA end was blocked by a Dig-anti-Dig F_ab_; this block ensures that synapsis occurs only at the fluorophore-labeled end.

**Figure 2:**
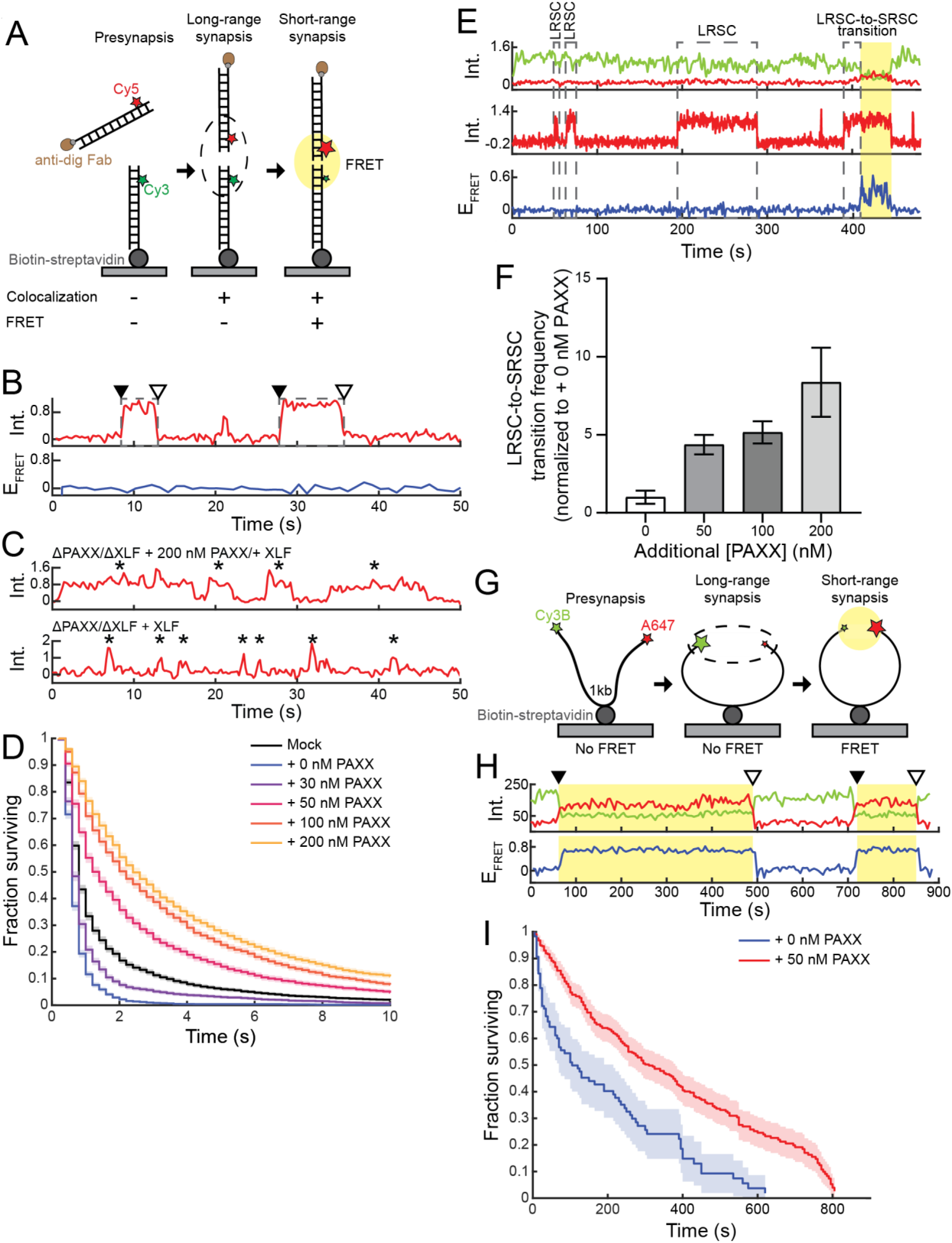
PAXX stabilizes both the LRSC and SRSC and facilitates the transition between the two synaptic complexes. (A) Schematic of two-piece substrate colocalization experiment. (B) Sample Cy5 intensity trace under Cy5 excitation (top) and FRET trace (bottom) in PAXX/XLF double depleted extract with 50 nM recombinant PAXX and 50 nM recombinant XLF rescue. FRET-free colocalization events indicative of LRSC formation are indicated with dotted boxes; the black arrow indicates the start of the colocalization event, and the white arrow indicates the end. (C) Sample Cy5 intensity traces under Cy5 excitation in PAXX/XLF double depleted extract with 50 nM recombinant XLF rescue and indicated recombinant PAXX rescue. Colocalization events indicated with asterisks. (D) Kaplan-Meier survival curves for Cy3-Cy5 colocalization events. All extracts, except for mock depleted extract, are PAXX/XLF double depleted extracts rescued with 50 nM recombinant XLF and indicated concentrations of recombinant PAXX. (E) Sample trace for data used to calculate synaptic transition efficiency showing donor (green) and acceptor (red) signals under donor excitation (top), acceptor signal under acceptor excitation (middle), and FRET efficiency (bottom). Trace shows LRSC events (without synaptic transition), indicated by acceptor signal under acceptor excitation without FRET, as well as LRSC-to-SRSC transition, in which an initial FRET-free state with positive acceptor signal is followed immediately by a FRET-positive state. LRSC-to-SRSC transition efficiency was calculated as the number of LRSC-to-SRSC transition events divided by the sum of the number of LRSC events and the number of LRSC-to-SRSC transition events. (F) Transition efficiency in undepleted egg extract supplemented with XRCC4/Lig4^K278R^ and indicated concentration of recombinant PAXX, normalized to the condition in which no recombinant PAXX was added. Error bars represent ± SD of bootstrapped data. (G) Schematic for one-piece substrate FRET experiment to measure SRSC stability. (H) Sample trace showing FRET events. FRET events are highlighted in yellow; start of FRET event is indicated by black arrow, and end of FRET event is indicated by white arrow. Traces show donor (green) and acceptor (red) signals under donor excitation (top) and FRET efficiency (bottom). (I) Kaplan-Meier survival curves for FRET events indicative of the SRSC. All extracts are PAXX/XLF/XRCC4 triple depleted extracts rescued with recombinant XLF, XRCC4/Lig4 ^K278R^, and any other indicated rescues. Shaded regions on Kaplan-Meier curves represent 95% confidence interval.

We previously showed that, upon addition of egg extract, the donor-and acceptor-labeled DNAs will colocalize upon formation of the LRSC but do not undergo Förster resonance energy transfer (FRET) (Figure 2A, B), as the distance between the two ends (>100 Å) is too great for FRET to occur. Transition to the SRSC brings the DNA ends into close proximity, which allows for FRET^8^ (Figure 2A). Acceptor-labeled DNAs readily colocalized with the surface-tethered DNAs even in the absence of PAXX (Figure 2C). From these trajectories, we calculated both the rate of formation of the LRSC and its stability. We measured the fraction of tethered DNA molecules that had experienced at least one colocalization event over time. These measurements showed comparable rates of association in mock depleted extract and in PAXX depleted extract rescued with varying concentrations of PAXX, suggesting that PAXX does not affect the formation rate of the LRSC (Extended Data 2A, B). In contrast, PAXX strongly stabilizes the LRSC once it forms. PAXX depletion reduced the mean LRSC dwell time by more than twofold relative to the LRSC dwell time in mock depleted extract (t_µ_ = 0.78 ± 0.02 s in PAXX depleted extract vs 1.67 ± 0.06 s in mock depleted extract) while supplementation of PAXX depleted extract with increasing PAXX concentrations progressively increased LRSC lifetime, plateauing at ∼3-fold above levels in mock depleted extract (t_µ_ = 4.69 ± 0.12 s at 200 nM PAXX) (Figure 2D).

We hypothesized that PAXX’s prolonging of LRSC lifetime promotes the LRSC-to-SRSC transition by providing time for the transition-enabling steps to occur. To test this hypothesis, we supplemented undepleted egg extract with varying concentrations of recombinant PAXX and performed the assay described in Figure 2A again, this time computing the fraction of FRET-free LRSC events that transitioned to a FRET-positive state that is indicative of the SRSC (Figure 2E). The fraction of LRSC events that transitioned to a high-FRET SRSC state increased with PAXX concentration, reaching an ∼8-fold enhancement at 200 nM PAXX relative to no added PAXX (Figure 2F). Thus, PAXX not only stabilizes the LRSC but also promotes its progression to the SRSC.

Finally, we sought to determine how PAXX impacts the stability of the SRSC. As the transition efficiency to the SRSC is low^8^ in the absence of excess PAXX, we used a fluorescently labeled intramolecular end joining substrate in which the ends are in high local concentration to increase the number of SRSC events we observed. This substrate consists of a 1 kb linear, blunt-ended DNA labeled with a donor fluorophore near one end and acceptor fluorophore near the other and contains an internal biotin for attachment to the flow cell surface (Figure 2G). As with the intermolecular tethering substrate, the appearance of a high-FRET state indicates transition to the SRSC (Figure 2G, H). Because the FRET efficiency of the short-range synaptic complex and the ligated product are indistinguishable, allowing ligation to occur would result in an overestimation of the SRSC lifetime^8^. As such, we prevented ligation by immunodepleting XRCC4/Lig4 from egg extract and rescuing with the catalytically inactive XRCC4/Lig4^K278R^. Importantly, this LIG4 mutant still supports the transition to the SRSC^8^. Rescuing PAXX-depleted extract with 50 nM PAXX nearly doubled the mean dwell time of the SRSC relative to the mean dwell time in depleted extract with no PAXX rescue (t_µ_ = 138.698 ± 17.369 s without PAXX rescue, t_µ_ = 262.414 ± 14.684 s with PAXX rescue) (Figure 2I), thus demonstrating that PAXX also stabilizes the SRSC, albeit more modestly than it stabilizes the LRSC.

### PAXX stabilizes XLF at the synaptic complex

Given that PAXX stabilizes synapsis, and that XLF and PAXX are genetically redundant^39–41^, we hypothesized that PAXX stabilizes XLF within the synaptic complex. To test this hypothesis, we carried out single-molecule imaging experiments to measure the stability of fluorescently labeled XLF and PAXX at DNA ends in egg extract. N-terminal ybbR-tags on XLF and PAXX were labeled with an acceptor fluorophore using Sfp synthase^47,48^. Both labeled proteins retained their activity and stimulated NHEJ comparably to their wild-type counterparts (Extended Data 2D-G).

To quantify protein stability within the synaptic complex, we measured the duration of colocalization of acceptor-labeled proteins with a 1000-bp donor-labeled circularizing DNA substrate identical to that used in Figure 2G except lacking an acceptor fluorophore (Figure 3). The lifetime of labeled XLF on DNA was measured in the presence or absence of unlabeled PAXX. We also performed the reciprocal experiment using labeled PAXX, measuring the effect of XLF on its stability. The association rates of XLF and PAXX were independent of the other protein (Extended Data 3A, B), and PAXX residence time was similar with or without XLF (t_µ_ = 5.52 ± 0.23 s in the presence of XLF, t_µ_ = 5.14 ± 0.21 without XLF; Figure 3A). In contrast, XLF was substantially more stable in the presence of PAXX (t_µ_ = 2.90 ± 0.24 s) than in its absence (t_µ_ = 1.75 ± 0.13 s) (Figure 3B).

**Figure 3:**
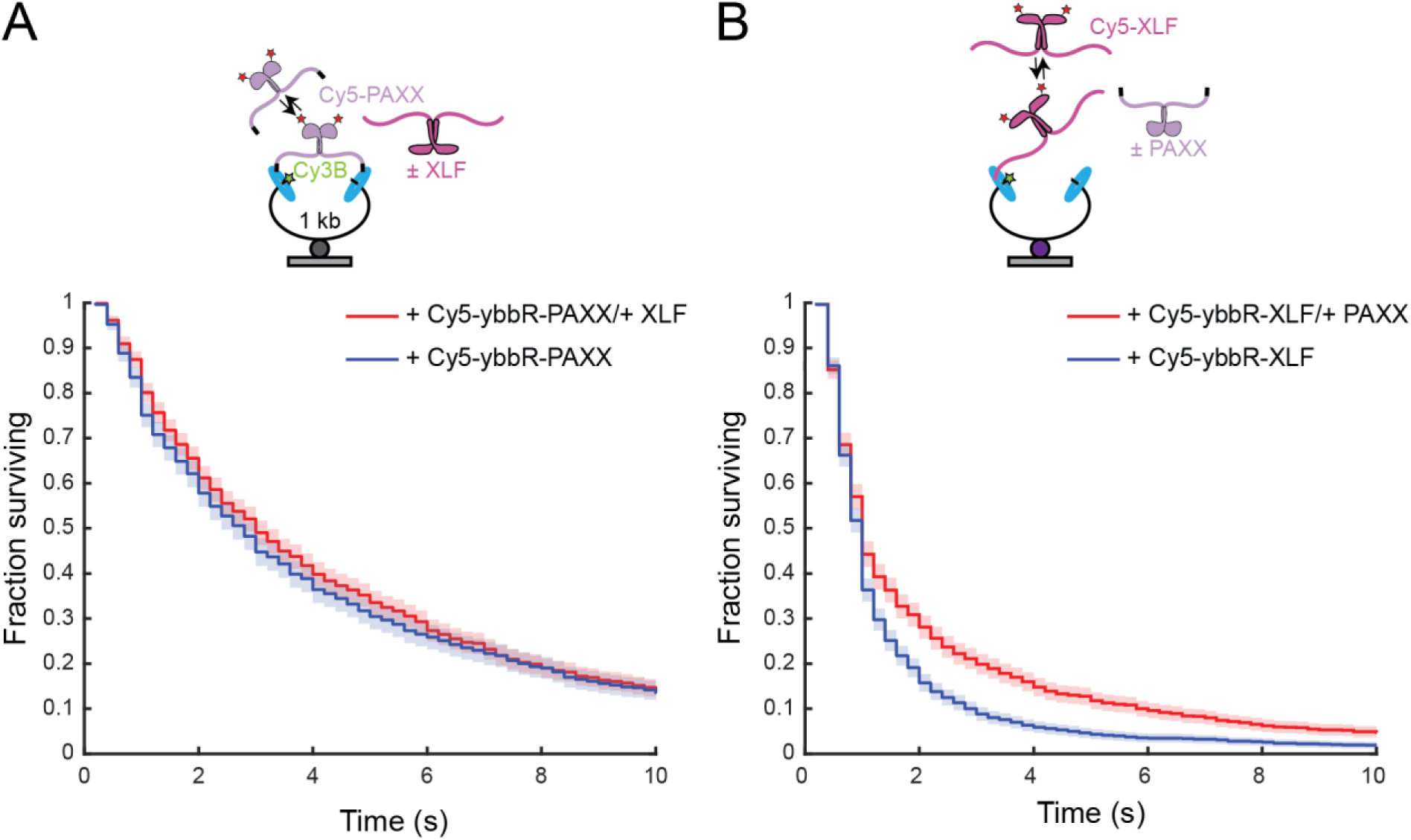
PAXX stabilizes XLF at the synaptic complex. (A) Kaplan-Meier curves showing stability of Cy5-PAXX on the pictured DNA substrate in the presence and absence of unlabeled XLF. (B) Kaplan-Meier curves showing stability of Cy5-XLF on the pictured DNA substrate in the presence and absence of unlabeled PAXX. Measurements were performed in PAXX/XLF double-depleted extract with the indicated rescues. Shaded regions on Kaplan-Meier curves represent 95% confidence interval.

We suggest this asymmetry reflects the stronger effect of PAXX on LRSC stability: prolonging the LRSC allows XLF to form additional stabilizing interactions within the complex, whereas the relatively weaker stabilization of the LRSC by XLF^8^ does not allow for similar stabilization of PAXX. Consistent with this model, XLF stability was unaffected by PAXX on a single-ended DNA substrate that cannot undergo synapsis (Extended Data 3C-D). Together, these results indicate that PAXX-dependent synapsis, particularly LRSC stabilization, stabilizes XLF, which in turn drives the LRSC-to-SRSC transition, providing a mechanistic basis for how PAXX facilitates SRSC formation.

### PAXX must bear at least one functional KBM and be of sufficient length to support NHEJ

As PAXX has no known enzymatic activity and no known binding partners other than Ku, its role in NHEJ is most likely a structural one. We hypothesize that PAXX stabilizes end synapsis by binding the Ku heterodimers present on both ends of the DSB using its tails, which tether the DNA ends together (Figure 4A). Indeed, structures of NHEJ synaptic complexes are consistent with a model in which PAXX bridges the DSB by binding Ku dimers on either end of the synaptic complex^23,49^; however, these structures do not preclude the possibility that this bridging interaction is not functionally necessary, as we have previously shown for XLF^10^. To test if this bridging interaction is necessary for PAXX’s stimulation of NHEJ, we generated several mutant PAXX constructs in which we altered the sequence or composition of various parts of its tails. As the KBMs of PAXX are essential to its stimulation of NHEJ in this model, we generated a PAXX construct bearing the mutation V198A/F200A (PAXX VF) (Figure 4B); these residues are well conserved, and the corresponding mutation in human PAXX disrupts the binding between human PAXX and Ku^20–23^. We then rescued PAXX-depleted extract with PAXX VF across a range of concentrations (∼10 nM to 1 µM) and assessed end joining in the rescued extracts at a single two-hour timepoint. Across all concentrations of PAXX VF tested, end joining was comparable to end joining in the absence of PAXX (Figure 4C-D).

**Figure 4:**
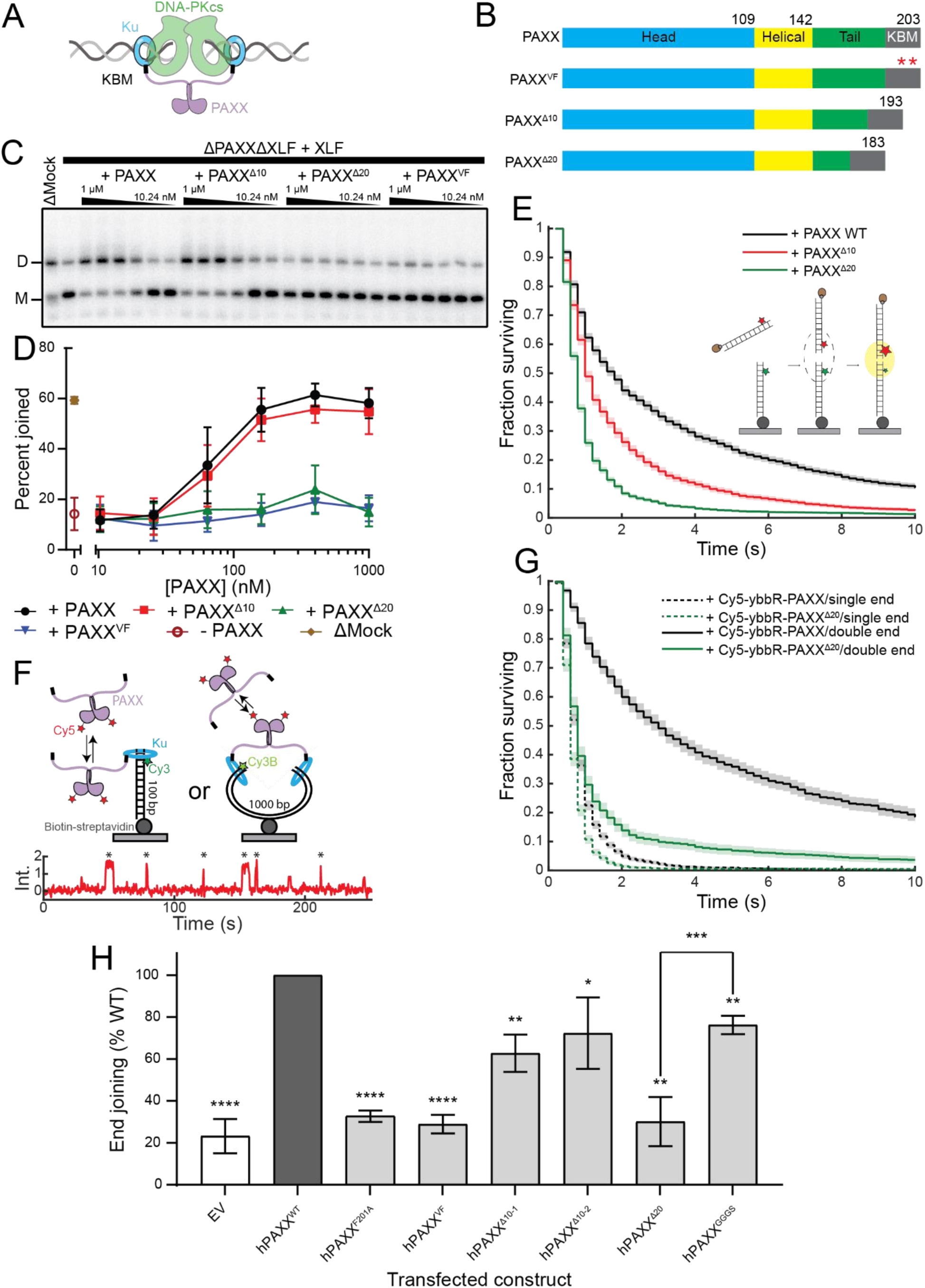
Shortening PAXX’s tails or deleting its KBMs impairs its stimulation of NHEJ. (A) Schematic of model for PAXX’s mechanism of action in NHEJ. (B) Shortened tail and KBM PAXX mutants. (C) End joining in extract rescued with indicated depletions and rescues. Each lane represents a two-hour timepoint. M: monomeric substrate, D: dimeric product. (D) Quantification of Figure 4C. Error bars represent ± SEM. Some information on depletion conditions has been omitted in the legend to save space. (E) Kaplan-Meier survival curves showing LRSC stability in PAXX/XLF double depleted extract rescued with recombinant XLF and indicated recombinant PAXX construct. (F) Schematic for fluorescence colocalization experiment to measure PAXX association to DNA ends and sample Cy5 trace (under Cy5 excitation) for experiment in Figure 4G. (G) Kaplan-Meier survival curves showing stability of Cy5-ybbR-PAXX and Cy5-ybbR-PAXX^Δ20^ at single-and double-ended DNA substrates. Extract used for this experiment was undepleted. Shaded regions in Figure 4E and Figure 4G represent 95% confidence interval. (H) End joining activity in human cells as measured by fluorescence reporter-based NHEJ assay in KO PAXX+XLF HEK-293T cells stably expressing the indicated human PAXX constructs EV: empty vector. Results are normalized to cells rescued with hPAXX^WT^ and plotted as mean values ± SD of 4**–**8 independent experiments (double KO and PAXX-WT: n = 8; all PAXX mutants: n = 4). Two-sided P-values from one-sample Student’s t-test between the PAXX-WT condition and the considered mutant are as follows and represented as asterisks: KO (<0.0001 ****), F201A (<0.0001 ****), VF (<0.0001 ****), Δ10-1 (0.0035 **), Δ10-2 (0.0475 *), Δ20 (0.0013 **), GGGS (0.0017 **). The two-sided P-value from unpaired two-sample Student’s t-test between the Δ20 and GGGS mutants was 0.0003 (***). * P < 0.05, ** P < 0.01, *** P < 0.001, **** P < 0.0001.

In addition to the requirement for functional KBMs, bridging requires that the tails of PAXX be long enough to span the DSB. We sought to test this hypothesis by generating PAXX mutants in which the tails were shortened by 10 and 20 residues (PAXX^Δ10^ and PAXX^Δ20^, respectively) (Figure 4B). Notably, to avoid perturbing the Ku binding motif^17,23^, we deleted residues from the N-terminal end of the PAXX tails. We selected these deletions based on distance measurements from structures of human PAXX (hPAXX) in complex with the LRSC. In particular, we measured the distance between the N-terminal residues of the resolved portions of hPAXX’s tails (C180) in two structures of hPAXX bound to different LRSCs^23^. For the “Ku mediated” LRSC (PDB: 8BH3), this distance is 131.322 Å, and for the “XLF mediated” LRSC (PDB: 8BHV), this distance is 103.488 Å^23^. We then estimated the distance (L) between the two C180 residues (or corresponding residues for shortened tail constructs) on the WT hPAXX homodimer and hPAXX homodimers in which the tails are shortened from the N-terminal side by 10 and 20 residues (hPAXX^Δ10^ and hPAXX^Δ20^, respectively), assuming that the tails are held taut and that each residue contributes 3.5 Å. Our estimations for L were 245 Å, 175 Å, and 105 Å for WT hPAXX, hPAXX^Δ10^, and hPAXX^Δ20^, respectively (Extended Data 4A). Consistent with the bridging model, L_WT_ is long enough to allow the tails of WT hPAXX to span the break in both LRSCs. L_Δ10_ is also long enough to allow hPAXX^Δ10^’s tails to span the break in both LRSCs; however, L_Δ20_ is too short to allow hPAXX^Δ20^’s tails to span the break in the “Ku mediated” LRSC. Though L_Δ20_ is barely long enough to allow for hPAXX^Δ20^’s tails to span the break in the “XLF mediated” LRSC, it would only be able to do so if the tails were held almost completely taut, which likely does not occur during synapsis. End joining in PAXX-depleted extract rescued with PAXX^Δ10^ was comparable to end joining in PAXX-depleted extract rescued with WT PAXX, while PAXX-depleted extract rescued with PAXX^Δ20^ was defective in end joining across all measured concentrations and comparable to PAXX-depleted extract rescued with PAXX VF or with no recombinant PAXX rescue (Figure 4C-D). Consistent with the bridging model, our results show that disrupting the interaction between PAXX and Ku or making deletions within PAXX’s tails that render the tails too short to span broken DNA ends both abolish PAXX’s ability to stimulate NHEJ. Single-molecule imaging using the intermolecular synapsis assay to measure LRSC stability in PAXX-depleted extract rescued with either WT PAXX or one of the shortened tail mutants further validates these results, as the LRSC becomes less stable as PAXX’s tail length decreases (WT PAXX: t_µ_ = 4.49 ± 0.15s, PAXX^Δ10^: t_µ_ = 2.15 ± 0.07 s, and PAXX^Δ20^: t_µ_ = 1.38 ± 0.06 s) (Figure 4E).

To ensure that the tail shortening mutations did not disrupt Ku binding and affect the ability of PAXX to localize to the DSB, we performed single-molecule imaging to directly observe the recruitment of PAXX to the synaptic complex. We bound donor-labeled DNA to a streptavidin-coated coverslip and introduced acceptor-labeled PAXX constructs (either WT or Δ20) in undepleted egg extract. We used both a 100 bp dsDNA substrate and a longer 1 kb circularizing substrate identical to the one presented in Figure 2G to measure PAXX association with both single DNA ends and the synaptic complex (Figure 4F). For both substrates, we observed dynamic colocalization of acceptor and donor signals, indicative of binding and unbinding of single PAXX molecules to the surface-bound DNAs (Figure 4F). The association rates and stabilities of both the WT and Δ20 PAXX constructs were comparable on the single end, indicating that the truncation mutation did not perturb the affinity of PAXX for Ku. However, on the circularizing substrate, WT PAXX associated more quickly and stably than PAXX^Δ20^, whose association rate and stability were comparable to those of either construct on the single-end substrate (Figure 4G and Extended Data 4E). This difference is likely a consequence of the inability of PAXX^Δ20^ to bind both Ku molecules simultaneously, which results in decreased avidity relative to WT PAXX.

To demonstrate that the defect in NHEJ stimulation that we observe in PAXX^Δ20^ is purely a result of shortening the PAXX tails and not due to loss of sequence-or composition-specific functions of the deleted residues, we generated two independent PAXX constructs in which the order of the first 20 N-terminal residues of the PAXX tail was randomly rearranged using the Sequence Manipulation Suite^50^, as well as a construct in which these first 20 residues were replaced with a (GGGS)_5_ linker (Extended Data 4B). Both constructs were able to rescue NHEJ in PAXX-depleted extract (Extended Data 4C). In addition, there is a serine that is present in PAXX^Δ10^ (S145, using the numbering for PAXX^Δ10^) that is not present in PAXX^Δ20^. To ensure that it is not a critical phosphorylation site whose deletion gives rise to the defect seen with PAXX^Δ20^, we generated a mutant in which S145 of PAXX^Δ10^ was mutated to Alanine (PAXX^Δ10^ S145A) (Extended Data 4B). While this mutant was slightly defective in NHEJ compared to WT PAXX, it was comparable to PAXX^Δ10^ (Extended Data 4C).

To extend our findings obtained in egg extract, we sought to determine the ability of shortened tail and KBM mutant PAXX constructs to stimulate NHEJ in human cells. To measure NHEJ in cells, we used a previously described fluorescence reporter assay^51^. Briefly, we transfected a Cas9-targeted reporter substrate into HEK-293T cells knocked-out for both PAXX and XLF (KO PAXX+XLF). Dual cleavage of this reporter substrate with Cas9 and subsequent error-free repair of the resulting DSB via NHEJ moves a GFP coding sequence in-frame and allows for the expression of functional GFP. We also integrated constructs in these cells for expressing several human hPAXX constructs analogous to the *X. laevis* constructs tested in egg extract. Notably, because the presence of XLF may mask the effects of PAXX, and because loss of XLF attenuates, but does not completely abolish, NHEJ in human cells^21,23^, we did not co-express XLF. Consistent with our results in egg extract, mutations to either F201 alone (Extended Data 5F) or both of V199 or F201 (analogous to V198 and F200 in *X. laevis* PAXX VF, respectively) in hPAXX (hPAXX^F201^ and hPAXX^VF^, respectively) resulted in a severe defect in NHEJ efficiency. Similarly, deleting the first 20 residues of hPAXX’s tails to generate hPAXX^Δ20^ also substantially attenuated NHEJ. In both cases, NHEJ efficiency was ∼30% of the efficiency obtained with WT hPAXX, with end joining in these cells comparable to end joining in cells transfected with empty vector controls (Figure 4H). In contrast, deleting the first or second 10 residues of the tails (hPAXX^Δ10-1^ and hPAXX^Δ10-2^, respectively) (Extended Data 5F) only modestly attenuated NHEJ (∼60% and ∼70% of NHEJ efficiency relative to hPAXX WT, respectively). In addition, we also introduced an hPAXX-expressing construct in these cells in which the first 20 N-terminal residues of the tail were replaced with a (GGGS)_5_ linker (hPAXX^GGGS^); consistent with our bridging hypothesis and our results in egg extract, this construct stimulated end joining at levels comparable to WT hPAXX (Figure 4H). All constructs tested were well-expressed in cells and localized to the nucleus, and hPAXX^Δ10-1^, hPAXX^Δ10-2^, and hPAXX^Δ20^ localized to sites of laser irradiation-induced DNA damage at comparable rates to WT hPAXX (Extended Data 5A-E, H). Our results demonstrate that, as in egg extract, PAXX’s stimulation of end joining in human cells requires both its ability to bind Ku and sufficiently long tails.

### PAXX requires two KBMs to facilitate NHEJ

To further test the bridging hypothesis, we asked whether both PAXX KBMs are necessary to stimulate NHEJ. We constructed well-defined tandem heterodimers of PAXX by linking the C-terminus of one PAXX monomer to the N-terminus of the other using a (GGGS)_3_ flexible linker. We then generated PAXX constructs bearing only a single functional KBM by mutating either the C-terminal or internal KBM to the non-Ku binding VF mutant (tdPAXX WT/VF and tdPAXX VF/WT, respectively). As a positive control, we also purified a tandem PAXX construct bearing both functional KBMs (tdPAXX WT/WT) (Figure 5A). We have successfully used a similar approach in the past to study the necessity of protein-protein interactions within XLF^10,14^. Using the experimental approach taken in Figure 4, we rescued PAXX-depleted extract with either wild-type PAXX or our tandem PAXX dimers across a range of concentrations and assessed end joining at a two-hour time point. tdPAXX WT/WT was able to robustly rescue NHEJ in PAXX-depleted extract, albeit in a slightly attenuated manner compared to WT PAXX. However, tdPAXX WT/VF and tdPAXX VF/WT were deficient in facilitating NHEJ compared to WT PAXX and tdPAXX WT/WT (Figure 5B-C). Taken together, these data support a model in which the PAXX KBMs do not serve merely to recruit PAXX to the synaptic complex but are also required for mediating the tethering of broken DNA ends via the PAXX tails.

**Figure 5:**
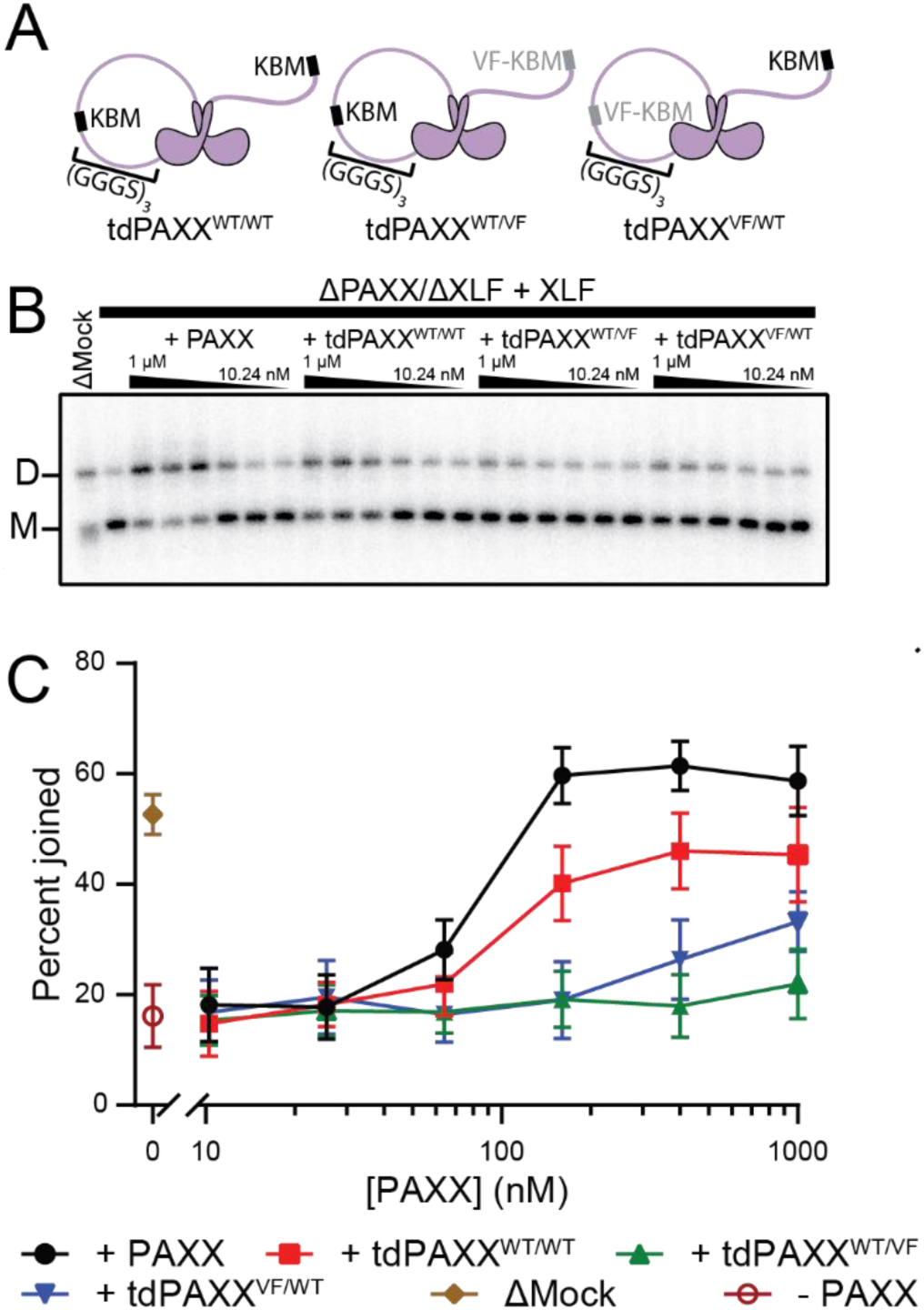
Each dimer of PAXX must bear two KBMs to stimulate NHEJ. (A) Tandem PAXX constructs. (B) End joining in extract with indicated depletion and rescues. Each lane represents a two-hour timepoint. M: monomeric substrate, D: dimeric product. (C) Quantification of Figure 5B. Some information about the depletion conditions has been omitted from the legend to save space. Error bars represent ± SEM.

While both tdPAXX WT/VF and tdPAXX VF/WT were deficient in rescuing PAXX depletion, tdPAXX VF/WT mediated a modest rescue at high concentrations compared to tdPAXX WT/VF, which was similarly deficient across all concentrations tested (Figure 5B, C). It is possible that there is a small amount of dimerization occurring between one coiled-coil domain of one tandem dimer and one coiled-coil domain of another; the resulting dimer of tandem dimers would contain two wild type KBMs and would thus be able to stabilize synapsis by bridging. Mass photometry measurements of the tandem dimer constructs, but not WT PAXX, reveal a small population of species with molecular weight that is approximately twice that of the tandem dimer (Extended Data 6); this is consistent with a minor population of dimers of tandem dimers in our protein preparations. While dimers of tandem dimers of both tdPAXX WT/VF and tdPAXX VF/WT would contain two wild type KBMs, we likely only observe the increase in rescue efficiency at high concentrations with tdPAXX VF/WT because its wild type KBMs are C-terminal rather than internal and are thus better able to bind Ku.

### PAXX and XLF have distinct roles during NHEJ

Our data thus far suggests a model in which the primary function of PAXX’s ordered domains is to dimerize its tails, which tether DNA ends to promote NHEJ. In contrast, our previous work has shown that XLF’s ordered domains, and particularly its head-to-head interaction with XRCC4, are the primary mediators of XLF’s function during NHEJ^10,14^. To test this model and to determine the relative contributions of each protein’s head and tail region, we generated a chimeric construct in which the tail domain of XLF was exchanged for the tail domain of PAXX (XLF^head/helix^-PAXX^tail^). As controls, we also generated the reciprocal chimeric construct bearing PAXX’s ordered domains and XLF’s tails (PAXX^head/helix^-XLF^tail^), as well as a variant of XLF^head/helix^-PAXX^tail^ in which point mutations that disrupt the essential interaction between XLF and XRCC4 (L68D/L117D)^10^ were introduced into the XLF head domains (XLF^head/helix^ ^LL→DD^-PAXX^tail^) (Figure 6A).

**Figure 6:**
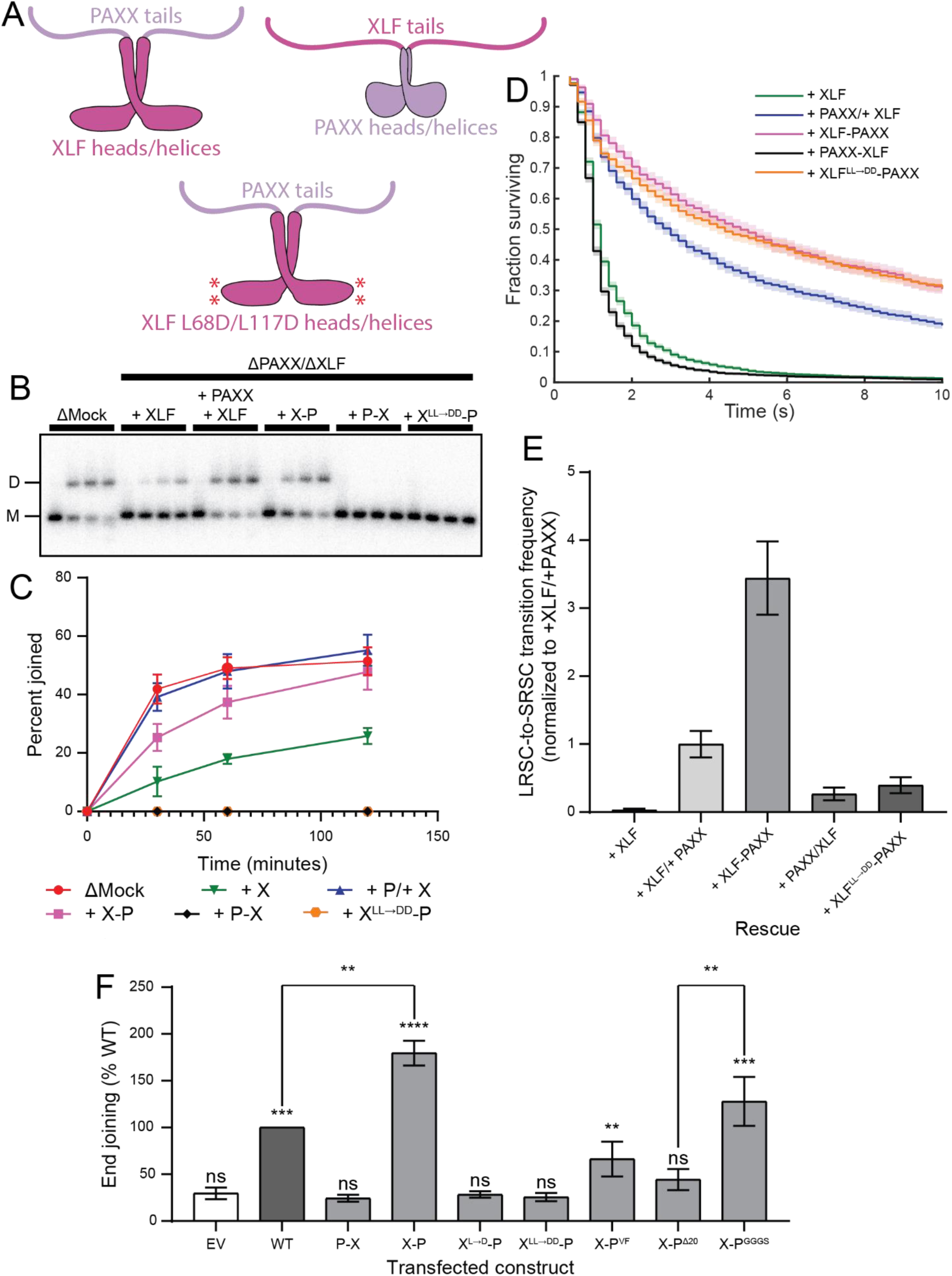
Chimeras of PAXX and XLF reveal complementary but distinct roles for different domains of each protein. (A) Domain architecture of *X. laevis* chimeric PAXX/XLF constructs. (B) End joining assay with indicated depletions and rescues to test chimeric PAXX constructs. Each construct was tested over a two-hour time course with samples withdrawn at 0, 30, 60, and 120 minutes. (C) Quantification of Figure 6B. Error bars represent ± SEM. Some information about the depletion conditions has been omitted from the legend to save space. (D) Stability of the long-range synaptic complex in PAXX/XLF double depleted extract with the indicated rescues. Shaded regions indicate 95% confidence interval. (E) Normalized LRSC-to-SRSC transition efficiency from experiment in Figure 6D. Transition efficiencies were calculated as the percentage of observed FRET-free LRSC events that exhibited transitions to a FRET-positive state indicative of the SRSC. Error bars represent ±SD of the bootstrapped data. (F) End joining as measured using the fluorescence reporter assay employed in Figure 4H in KO PAXX+XLF stably expressing the indicated recombinant constructs. EV: empty vector. WT: WT hPAXX. P-X: hPAXX^head/helix^-hXLF^tail^. X-P: hXLF^head/helix^-hPAXX^tail^. X^L→D^-P: hXLF^head/helix^ ^L115D^-hPAXX^tail^. X^LL→DD^-P: hXLF^head/helix^ ^L65D/L115D^-hPAXX^tail^. X-P^VF^: hXLF^head/helix^-hPAXX^tail^ with VF mutation in PAXX tail. X-P^Δ20^: hXLF^head/helix^-hPAXX^tail^ with 20 residue internal deletion in PAXX’s tail. X-P^GGGS^: hXLF^head/helix^-hPAXX^tail^ ^GGGS^. Two-sided P-values from unpaired two-sample Student’s t-test (one-sample test for the KO vs PAXX-WT comparison) between the double KO and the considered condition are as follows and represented as black asterisks: hPAXX-WT (0.0002 ***), P-X (0.1863 ns), X-P (<0.0001 ****), X^L→D^-P (0.7272 ns), X^LL→DD^-P^WT^ (0.3180 ns), X-P^VF^ (0.0099 **), X-P^Δ20^ (0.0637 ns), X-P^+20^ (0.0003 ***). The two-sided P-values from one-sample Student’s t-test between hPAXX-WT and X-P and unpaired two-sample Student’s t-tests between X-P^Δ20^ and X-P^+20^ are 0.0012 (**) and 0.0011 (**), respectively. ** P < 0.01, *** P < 0.001, **** P < 0.0001, ns: not significant.

Individual chimeras were added to extract depleted of both PAXX and XLF to determine their ability to facilitate end joining. Of the three chimeras, only XLF^head/helix^-PAXX^tail^ was able to robustly rescue NHEJ, but not PAXX^head/helix^-XLF^tail^ or XLF^head/helix^ ^LL→DD^-PAXX^tail^, where little to no end joining was observed (Figure 6B, C). While previous work has shown that mutations within XLF’s KBM abolish NHEJ^14^, our results suggest that XLF’s KBM only functions to recruit XLF to DSBs and can be substituted with other sequences that also have affinity for Ku. Therefore, these results demonstrate that only XLF’s ordered domains and PAXX’s tail are necessary and sufficient for the functions of both PAXX and XLF in NHEJ.

To further determine specific roles for XLF’s ordered domains and PAXX’s tails, we performed single-molecule imaging using the assay described in Figure 2A. Once again, we rescued PAXX/XLF double depleted extract with either the chimeric constructs or wild type PAXX or XLF. Consistent with our result that PAXX does not affect the formation rate of the LRSC (Extended Data 2A-B), the formation rate of the LRSC was similar across all conditions (Extended Data 2C). Both XLF^head/helix^-PAXX^tail^ and XLF^head/helix^ ^LL→DD^-PAXX^tail^ stabilized long range synapsis; however, PAXX^head/helix^-XLF^tail^ did not (Figure 6D). Given that XLF^head/helix^ ^LL→DD^-PAXX^tail^ stabilizes long range synapsis but cannot promote NHEJ, its defect is likely that it cannot facilitate transition to the SRSC. We thus conclude from these results that PAXX’s stimulation of NHEJ primarily arises from stabilization of synapsis via its tails, with its ordered domains only serving to dimerize the tails, whereas XLF’s ordered domains and their interaction with XRCC4 directly mediates long-to-short range synaptic transition. Consistent with this model, LRSC-to-SRSC transition efficiency is more than threefold greater in PAXX/XLF depleted extract rescued with XLF^head/helix^-PAXX^tail^ than in depleted extract rescued with both XLF and PAXX (Figure 6E), as the chimeric construct eliminates the need for XLF to be recruited separately from PAXX, reducing the number of steps required to assemble a transition-competent LRSC. Therefore, PAXX and XLF serve complementary but distinct roles in synapsis.

As before, we validated these results in human cells using the approach described in Figure 4H. Consistent with our data in egg extract, end joining in KO PAXX+XLF cells stably expressing a construct bearing hPAXX’s head and helical domains and XLF’s tail domains (hPAXX^head/helix^-hXLF^tail^) was deficient in NHEJ (∼25% efficiency relative to hPAXX WT) and comparable to the empty vector control. In contrast, the reciprocal construct (hXLF^head/helix^-hPAXX^tail^) stimulated NHEJ to an even greater extent than hPAXX on its own (∼175% relative to hPAXX WT), likely due to the simultaneous contributions of PAXX’s tails and XLF’s heads. The human analogues of XLF^head/helix^ ^LL→DD^-PAXX^tail^, carrying mutations in one or both lysines in the XLF head domain that contact XRCC4 were similarly deficient (Figure 6F). Chimeras bearing XLF’s ordered domains and PAXX’s tails bearing either the VF non-Ku binding mutation or a 20-residue internal deletion (Extended Data 5G) stimulated end joining to an intermediate level between full XLF/PAXX double KO and rescue with WT PAXX. Finally, consistent with our results in Extended Data 4C, replacing the first 20 residues of the PAXX tail on hXLF^head/helix^-hPAXX^tail^ with a (GGGS)_5_ linker (hXLF^head/helix^-hPAXX^tail^ ^GGGS^) (Extended Data 5G) did not alter its ability to stimulate NHEJ relative to hXLF^head/helix^-hPAXX^tail^ (Figure 6F). These results suggest that the complementary yet distinct roles of PAXX tails and XLF heads are conserved between *Xenopus* egg extract and human cells.

## Discussion

Since its discovery, the mechanistic role of PAXX in NHEJ and the basis for its genetic redundancy with XLF have remained unclear. Here, we show that PAXX stabilizes synapsis, particularly the early stages of the LRSC in which the “Ku-mediated” LRSC forms, by using its disordered tails to bridge DNA ends through KBM-dependent binding of Ku. This synaptic stabilization increases the residence time of XLF at DNA ends, allowing for efficient formation of the “XLF-mediated” LRSC and subsequent LRSC-to-SRSC transition and ligation (Figure 7A). Using tail and KBM mutants and domain-swapped chimeras, we further separate the core activities of PAXX and XLF in NHEJ. Dimerized PAXX tails are sufficient to stabilize synapsis, whereas XLF’s ordered domains are required to drive the LRSC-to-SRSC transition via interaction with XRCC4.

**Figure 7:**
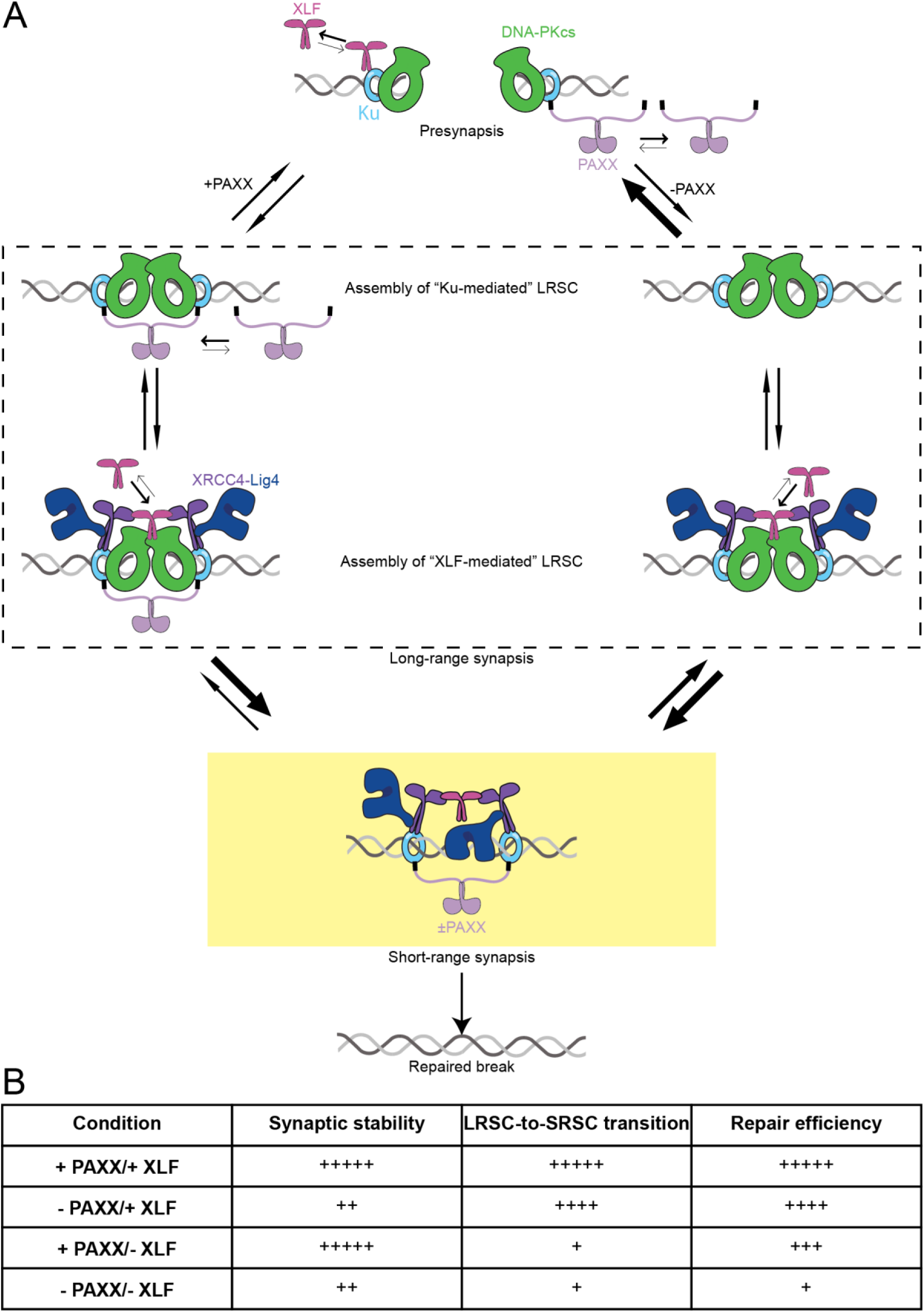
A mechanistic model explaining PAXX’s role in NHEJ. (A) PAXX stabilizes all stages of synapsis by tethering broken DNA ends together using its tail domains and their interaction with Ku in a manner independent of the identity of the domain that dimerizes the tail domains. PAXX’s stabilization of the “Ku-mediated” LRSC during the early stages of synapsis allows for efficient assembly of downstream NHEJ factors such as XLF, which are more stable during synapsis than during presynapsis and are required for forming the “XLF-mediated” LRSC, which is the direct precursor to the SRSC.The absence of PAXX destabilizes synapsis and, as a result, the binding of downstream NHEJ factors to the synaptic complex. (B) PAXX’s stabilization of synapsis and XLF’s stimulation of synaptic transition are distinct but complementary roles. Defects in one or the other result in a slight but tolerable defect in NHEJ that still allows for viability, but defects in both result in a cumulative and substantial overall defect in NHEJ and embryonic lethality.

Structural studies have shown that PAXX can bridge a DSB by simultaneously engaging Ku molecules on opposing ends through its C-terminal KBMs^23,49^. However, because substantial portions of PAXX, including much of its tails and ordered domains, are unresolved in these structures, it is unclear whether bridging is functionally required or whether the tails primarily act to recruit PAXX to Ku-bound ends. Our mutational analysis directly supports a bridging mechanism, as disruption of the KBM–Ku interaction, shortening the tail below a critical length, or enforcing a PAXX dimer with only one functional KBM strongly diminishes PAXX-dependent stimulation of end joining.

Our single-molecule assays further define when PAXX acts during NHEJ. PAXX stabilizes both long-and short-range synaptic complexes, with a substantially more pronounced effect on LRSC lifetime than on SRSC lifetime. In contrast, we do not detect a major effect of PAXX on the formation rate of the LRSC, suggesting that PAXX primarily acts after initial end capture to suppress premature dissociation of synaptic intermediates. This kinetic role helps reconcile why PAXX can have modest phenotypes in some genetic contexts yet become important under conditions that challenge end capture and retention. Consistent with this idea, we observe a larger dependence on PAXX for intermolecular joining than for intramolecular joining substrates, where high local end concentration partially bypasses a requirement for synaptic stabilization.

Our results also provide a mechanistic explanation for the widely observed genetic interaction between PAXX and XLF. In mice, single knockouts of PAXX or XLF cause relatively mild phenotypes, whereas combined loss results in embryonic lethality^39–41^. Accordingly, it has been proposed that PAXX and XLF perform overlapping functions in parallel NHEJ pathways^23^. However, prior biochemical, cellular, and structural studies have suggested non-equivalent roles: PAXX loss produces smaller defects than XLF loss in reconstituted systems^15–17^, XLF-deficient cells are often more sensitive to genotoxic stress than PAXX-deficient cells^21,23,52^ and PAXX and XLF bind Ku independently at distinct sites^23^. Our data support a sequential division of labor in which PAXX primarily stabilizes the LRSC, thereby increasing the likelihood that XLF can productively engage XRCC4/Lig4 to drive LRSC-to-SRSC transition. In this view, “redundancy” arises not because PAXX and XLF execute identical steps in parallel, but because loss of either one produces a partial defect—destabilization of the early stages of synapsis (PAXX) or impaired synaptic maturation (XLF)—that can be buffered by remaining pathway capacity, whereas combined loss compromises both stability and transition and yields a severe NHEJ defect (Figure 7B). This framework also helps rationalize genetic interactions among additional NHEJ factors. Like PAXX, DNA-PKcs stabilizes the LRSC^6,8,12^ and is synthetically lethal with XLF^53–55^, consistent with two defects that jointly impair synaptic stabilization and synaptic maturation. In contrast, PAXX deletion does not exacerbate loss of DNA-PKcs^54^, consistent with PAXX and DNA-PKcs contributing overlapping synapsis-stabilizing functions. Conversely, combining PAXX deletion with inhibition of DNA-PKcs catalytic activity, which has been implicated in synaptic complex transition^8,56–58^, produces a stronger defect than either perturbation alone^59^. Together, these results support a general principle for NHEJ genetics that factors that stabilize synapsis can show synthetic interactions with factors that promote synaptic transition, even when their molecular activities are distinct; such a framework might explain other observed genetic redundancies, such as the redundancy observed between XLF and either MRI/CYREN^59,60^ or ATM^61^.

Our finding that synapsis stabilization can be conferred by dimerized PAXX tails raises an evolutionary question: why does vertebrate PAXX retain XRCC4-family head and helical domains? One possibility is suggested by XRCC4-family diversification across eukaryotes. A PAXX homolog in *Arabidopsis thaliana* retains Ku binding through its tails but also exhibits XLF-like interactions with XRCC4 via its head domains^62^, raising the possibility that ancestral XRCC4-family proteins combined tethering and transition-promoting activities that later partitioned into distinct vertebrate paralogs. Alternatively, PAXX head domains may contribute to regulation, localization, or context-specific partner interactions *in vivo* that are not captured by the substrates and conditions assayed here. Indeed, a prior study reported PAXX head-domain mutations attenuate NHEJ^21^, but this effect is likely attributable to poor expression of the mutant protein^60^.

While this work defines the mechanistic basis for PAXX’s stimulation of NHEJ, it also raises several questions. First, our experiments primarily use blunt, ligation-compatible ends and thus do not address how PAXX contributes to repair of ends that require processing. Recent SRSC structures containing DNA polymerase µ (Pol µ) suggest that access to DNA ends by processing factors may require transient rearrangements of the ligation machinery^49^, which could further destabilize the SRSC and thereby increase reliance on PAXX-mediated stabilization under processing-required conditions. Consistent with this idea, biochemical reconstitutions have suggested a larger requirement for PAXX when Pol λ–dependent end processing precedes ligation^16^. Applying the single-molecule assays used here to a panel of end chemistries would help define how synaptic stabilization and synaptic maturation are tuned to different substrates.

Second, our chimera experiments suggest that XLF may contribute additional contacts within the synaptic machinery beyond the canonical head-to-head interaction with XRCC4. Despite disrupting XRCC4 binding, the XLF^head/helix^ ^LL→DD^-PAXX^tail^ chimera stabilizes the LRSC slightly more than WT PAXX and XLF (Figure 6D) yet fails to support end joining in egg extract (Figure 6B-C) and in cells (Figure 6F). One possibility is that, as suggested by previous structural work, XLF’s coiled-coil or C-terminal regions engage Ku or other synaptic components^31,49^, allowing the chimera to occupy space within the synaptic complex while failing to recruit or stabilize XRCC4/Lig4 in a productive configuration. Defining these additional XLF contacts, their strength, and their dependence on synaptic state will be important for a complete mechanistic description of synaptic maturation.

In sum, our results support a model in which NHEJ fidelity and efficiency emerge from a coordinated sequence of modular activities: Ku-anchored tethers such as PAXX (and DNA-PKcs) stabilize the early stages of end synapsis, thereby creating a kinetic window for assembly of maturation factors such as XLF–XRCC4/Lig4 that drive formation of the ligation-competent SRSC. This sequential organization explains how factors can appear genetically redundant while remaining mechanistically distinct and provides a framework for understanding how NHEJ adapts to diverse end structures and repair contexts.

## Data tables for single molecule imaging experiments

All errors represent ±SD of the bootstrapped data (1000 rounds of bootstrapping)

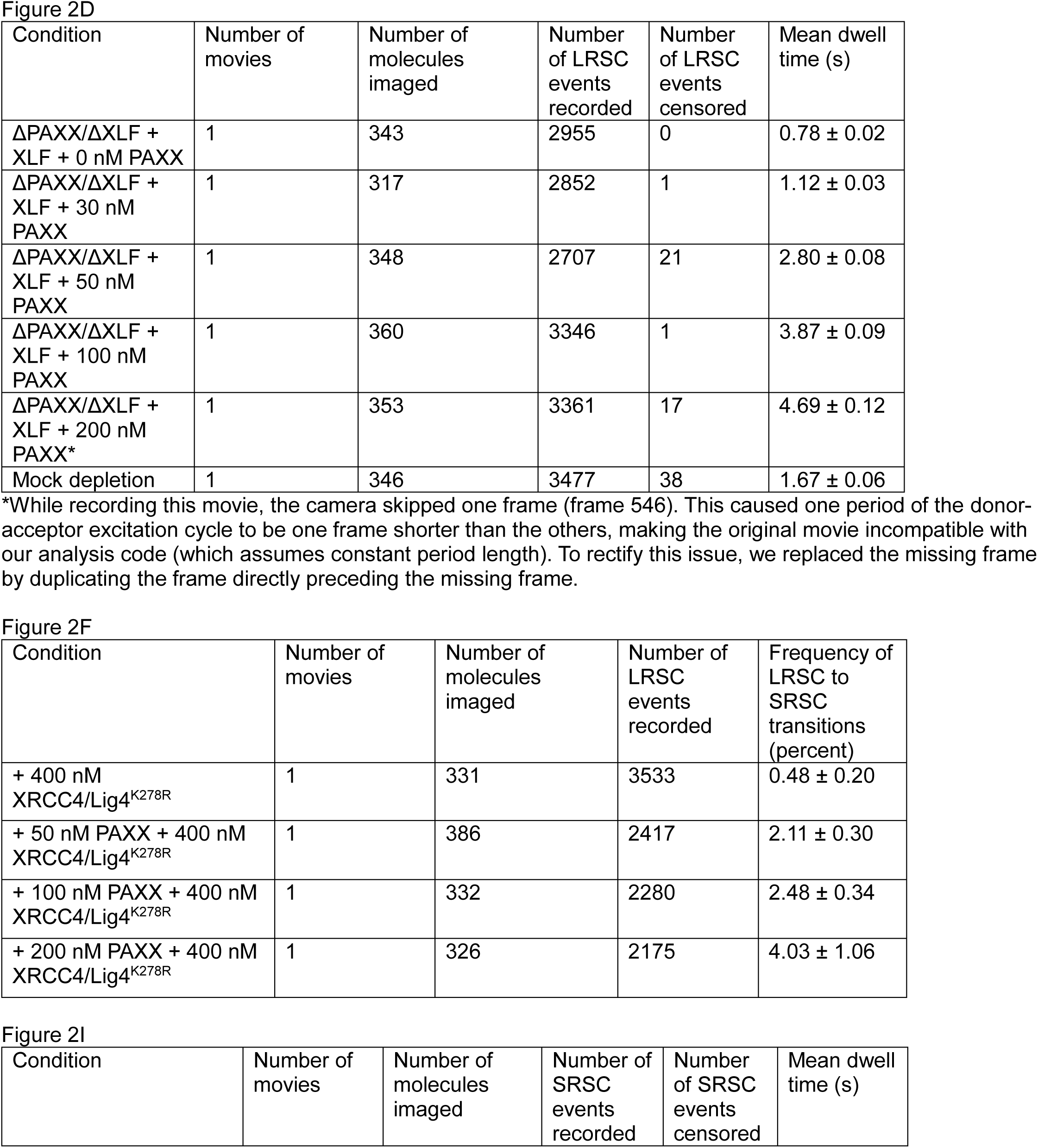

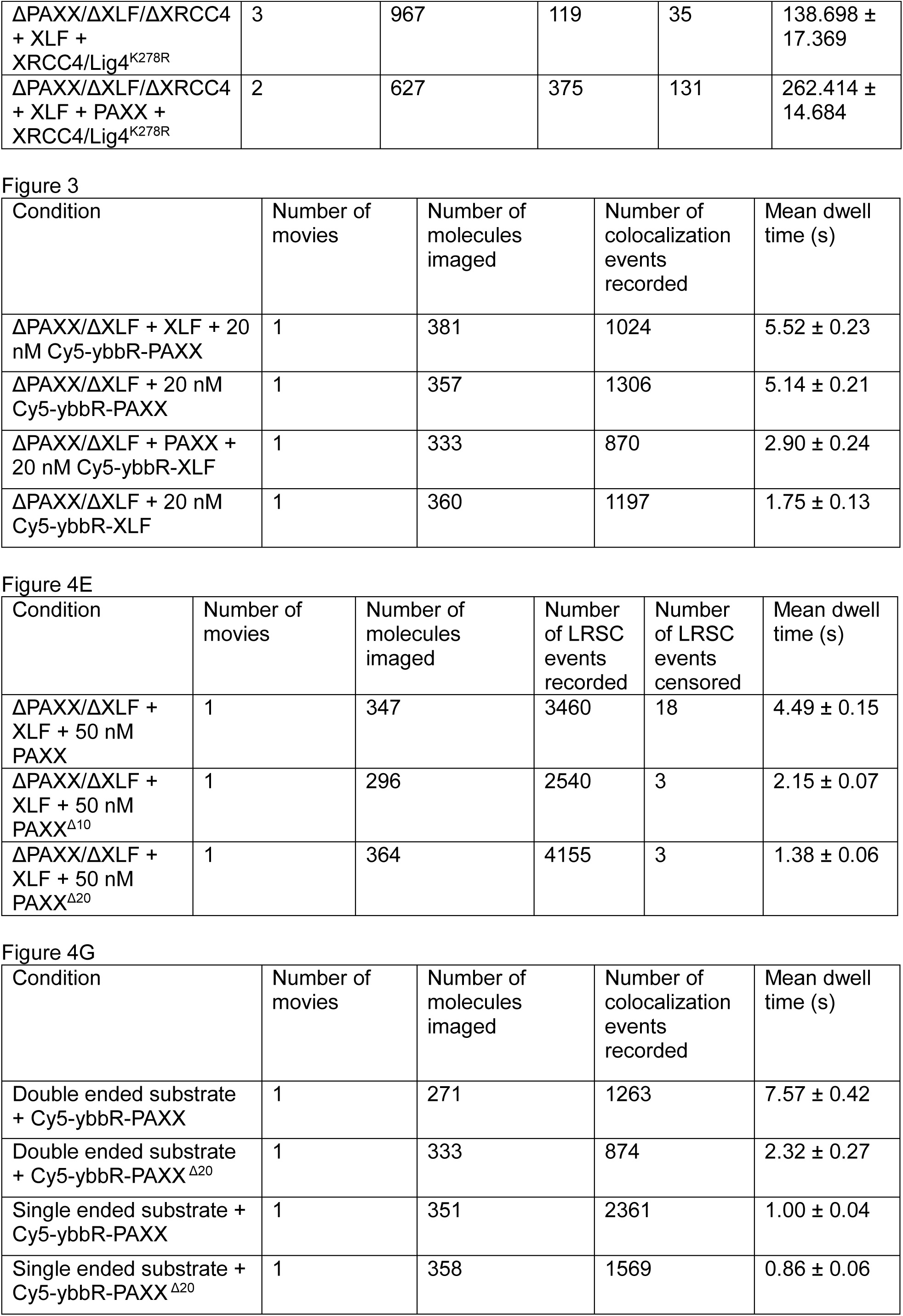

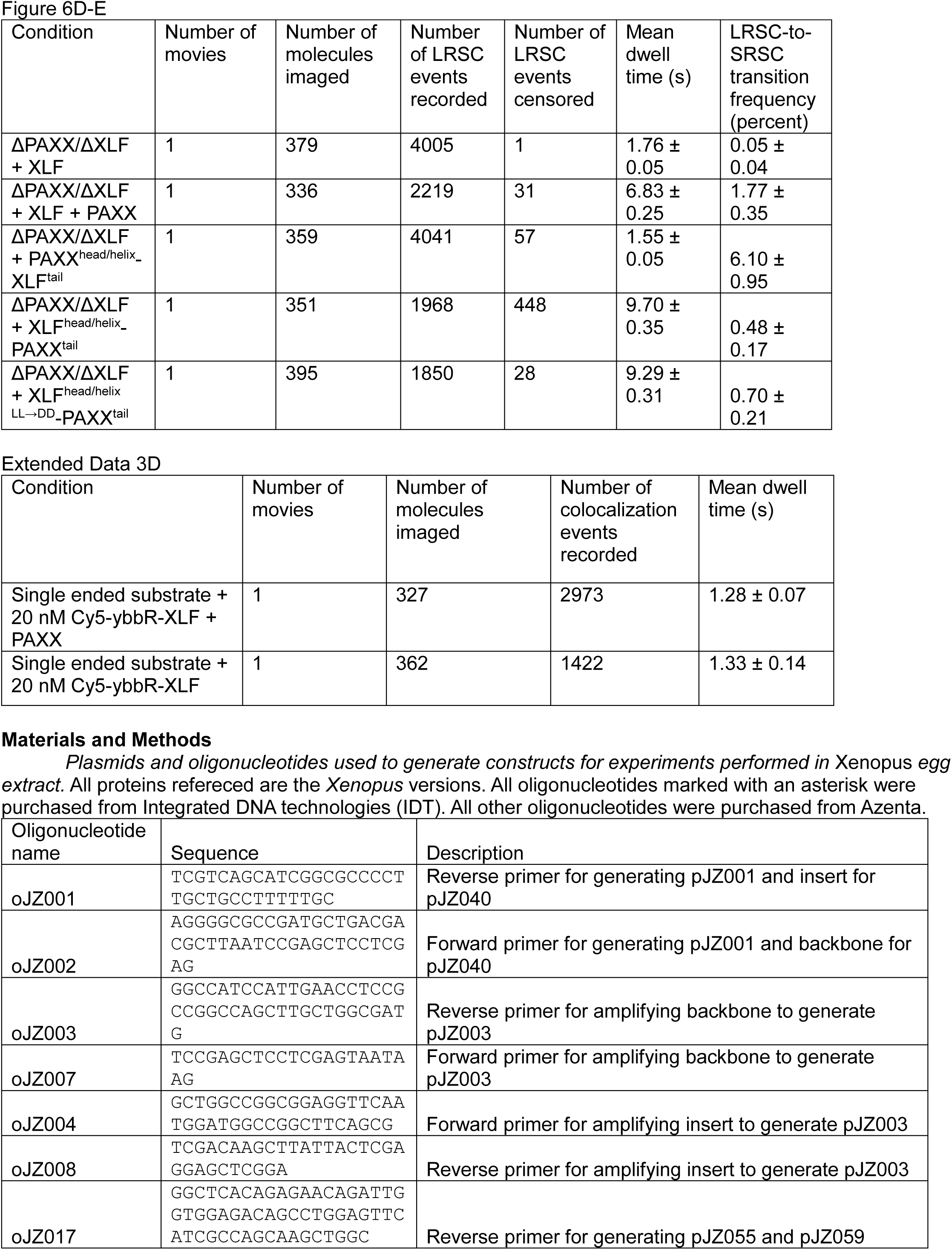

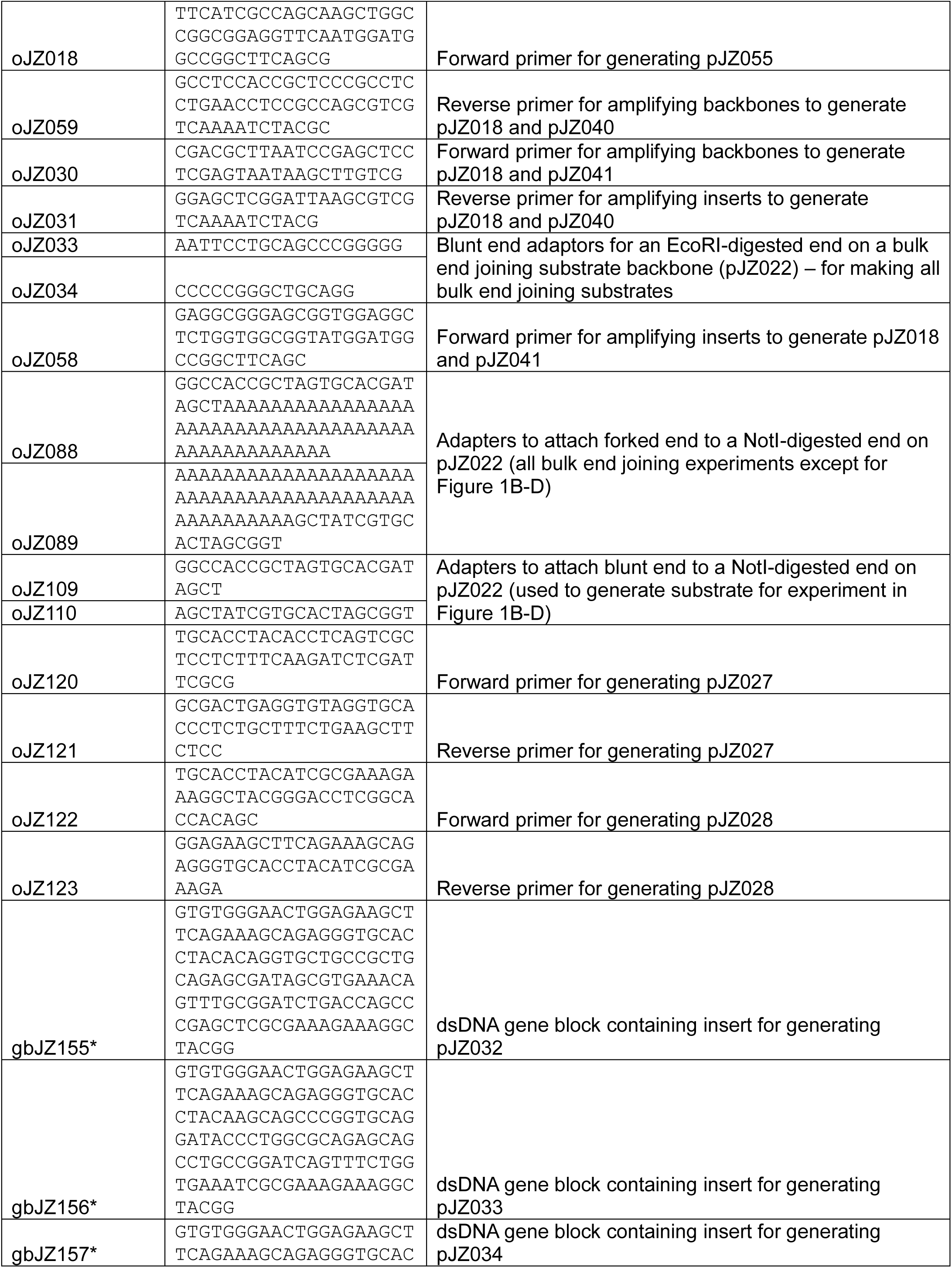

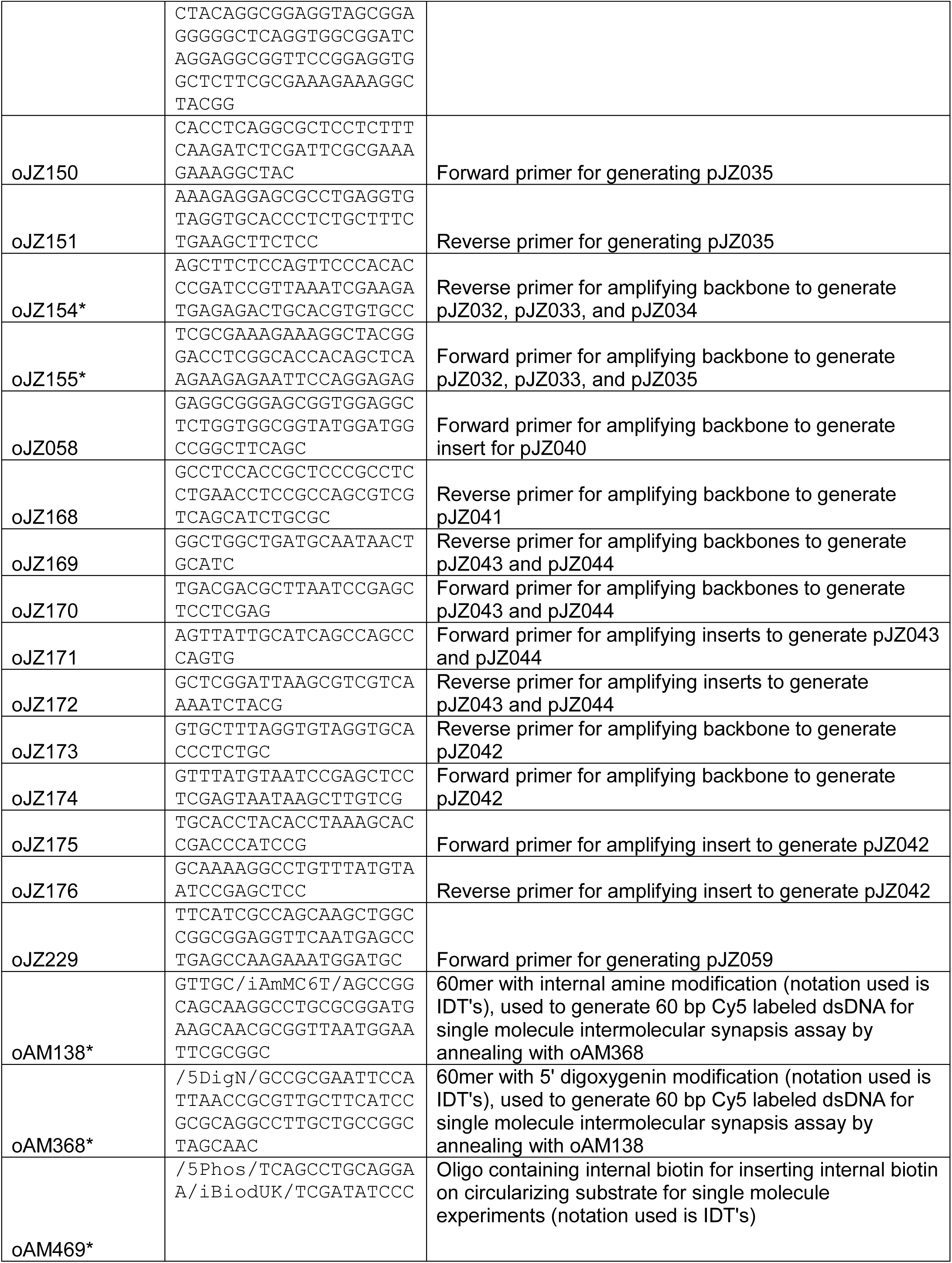

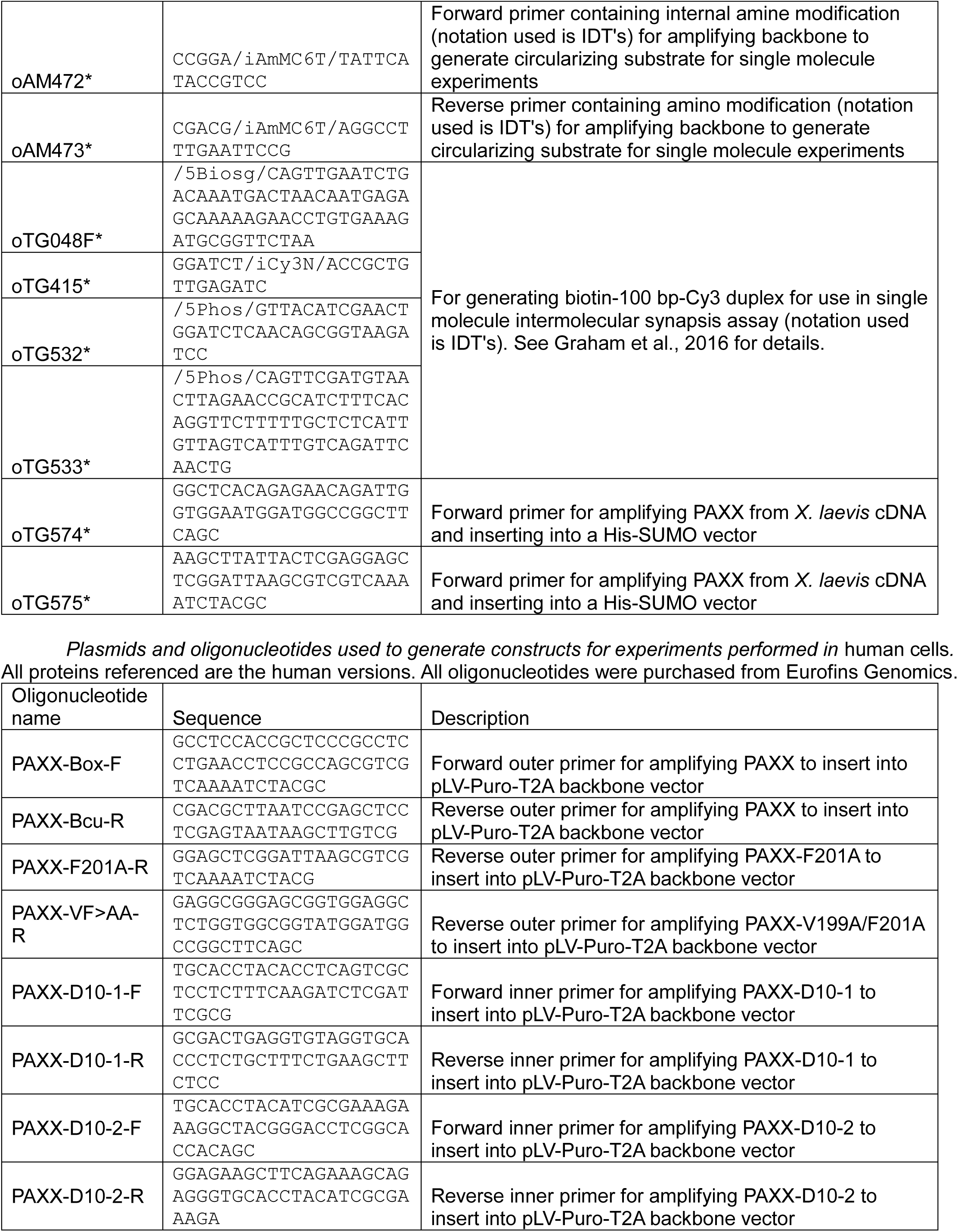

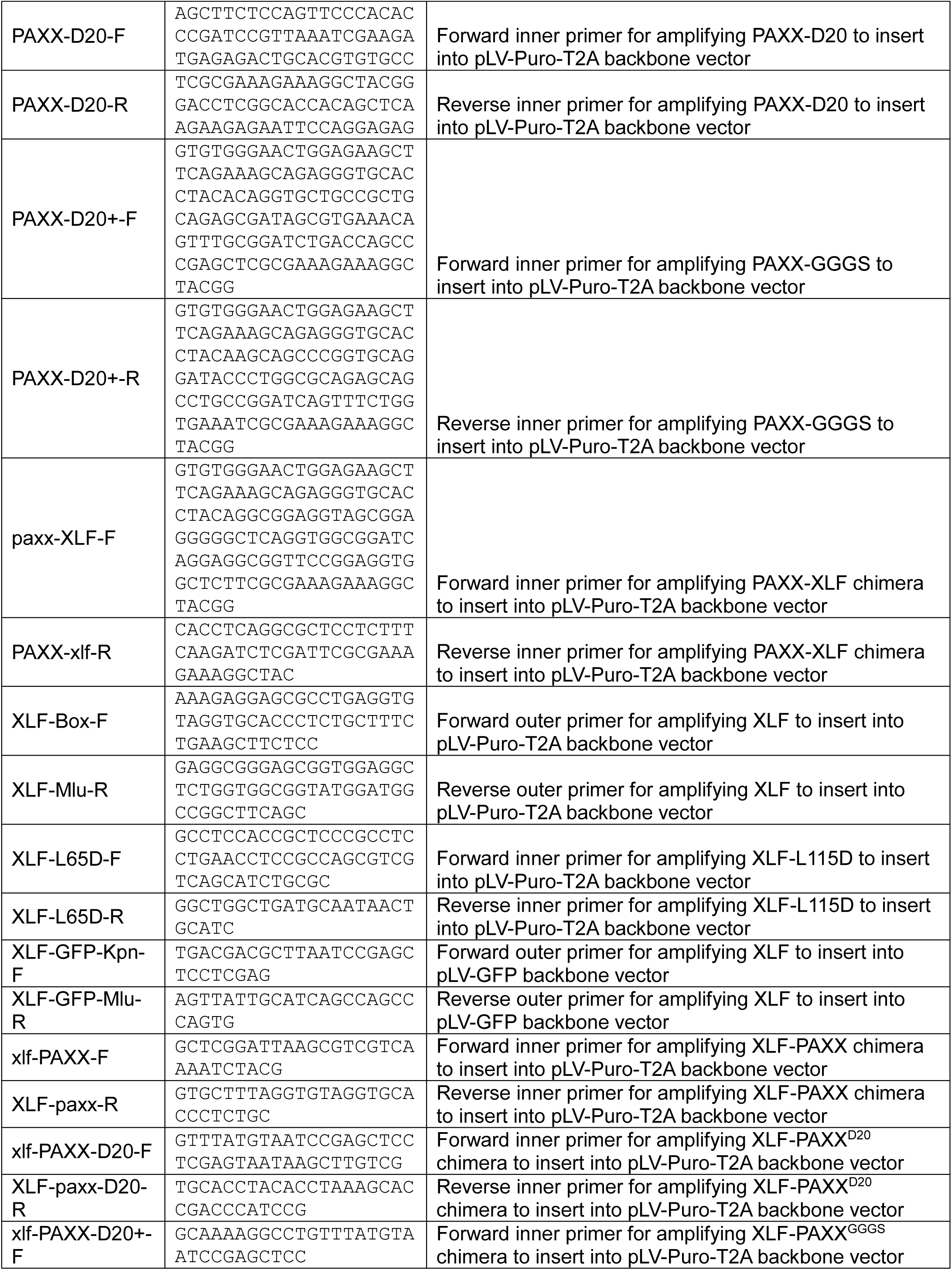

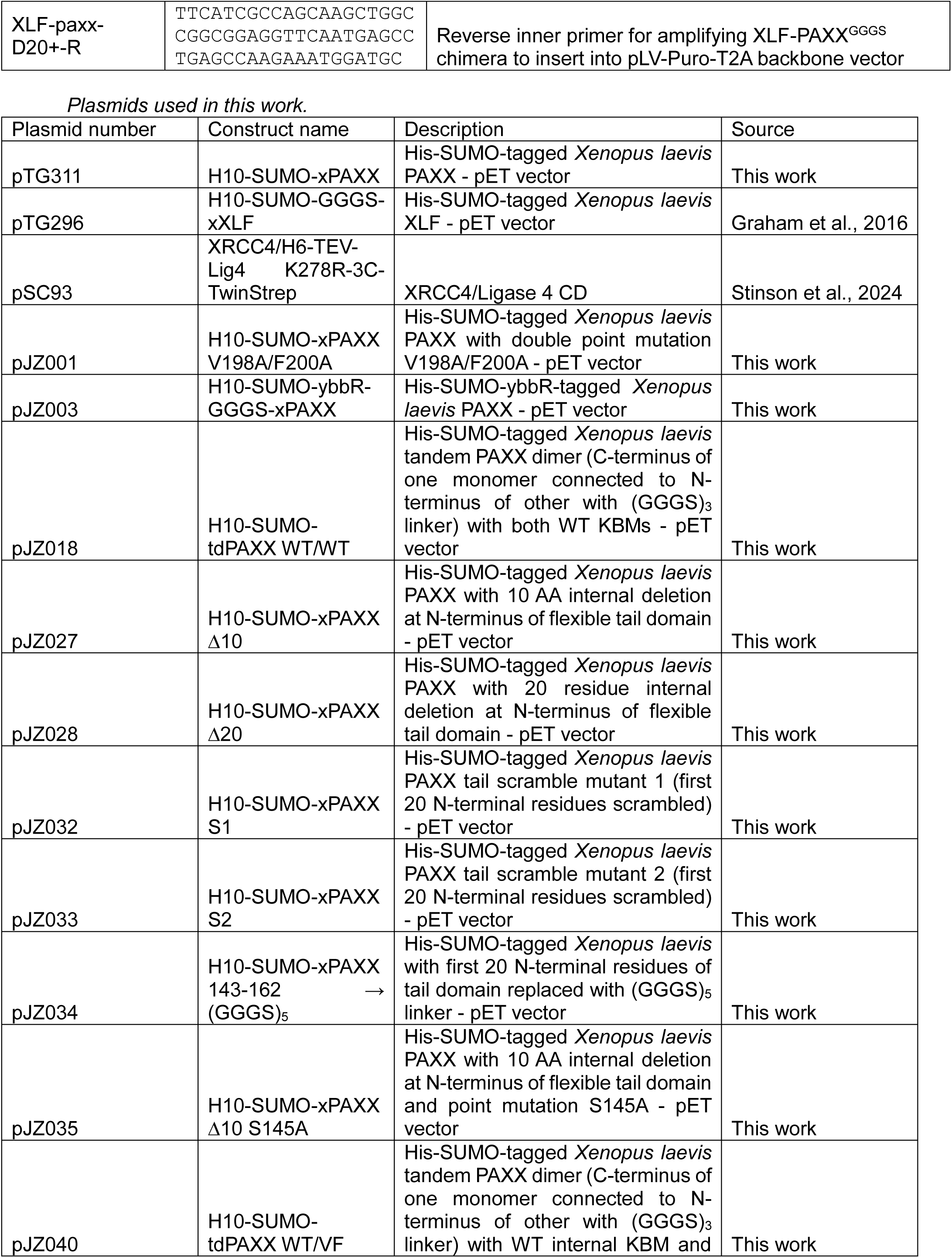

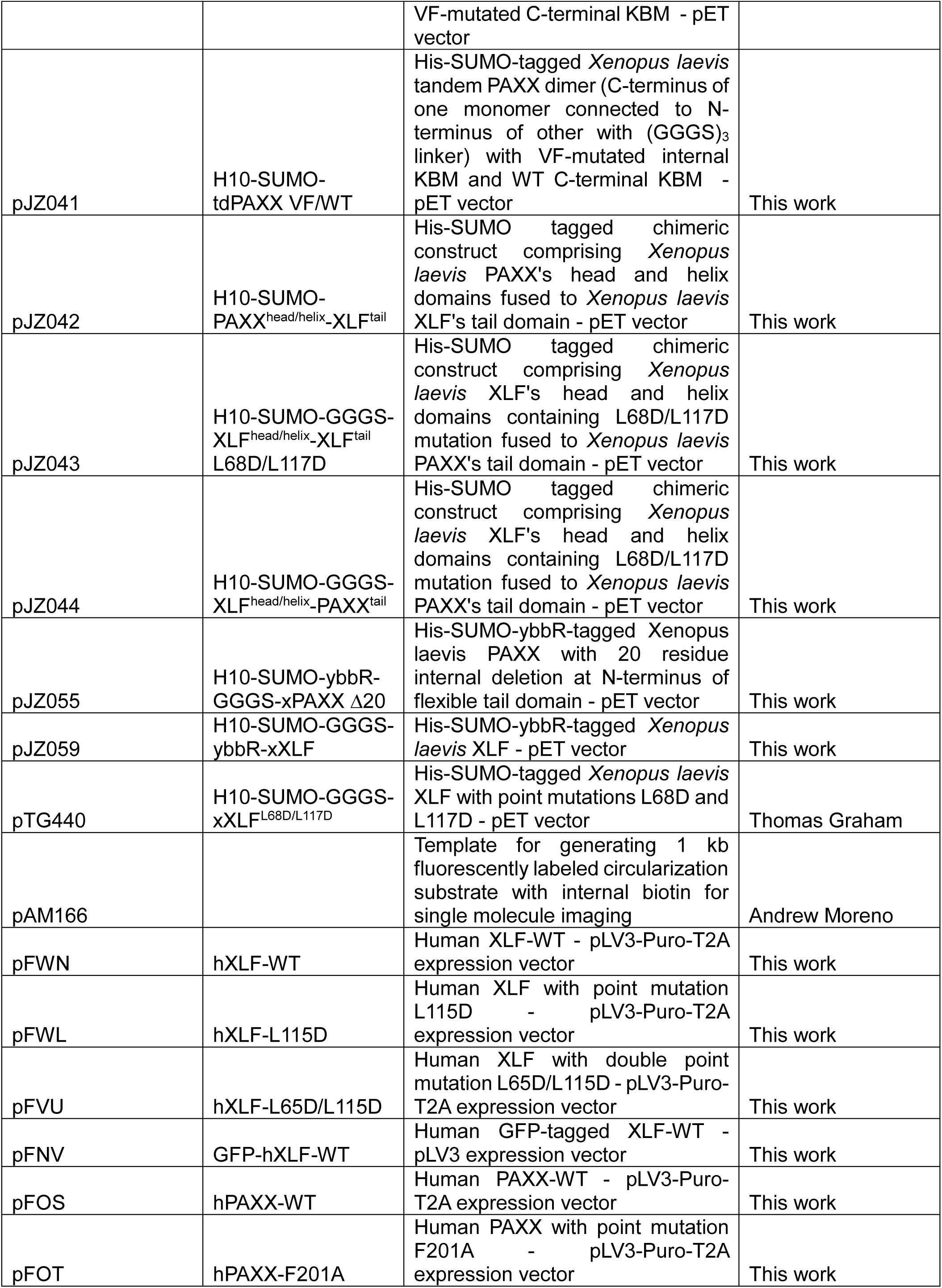

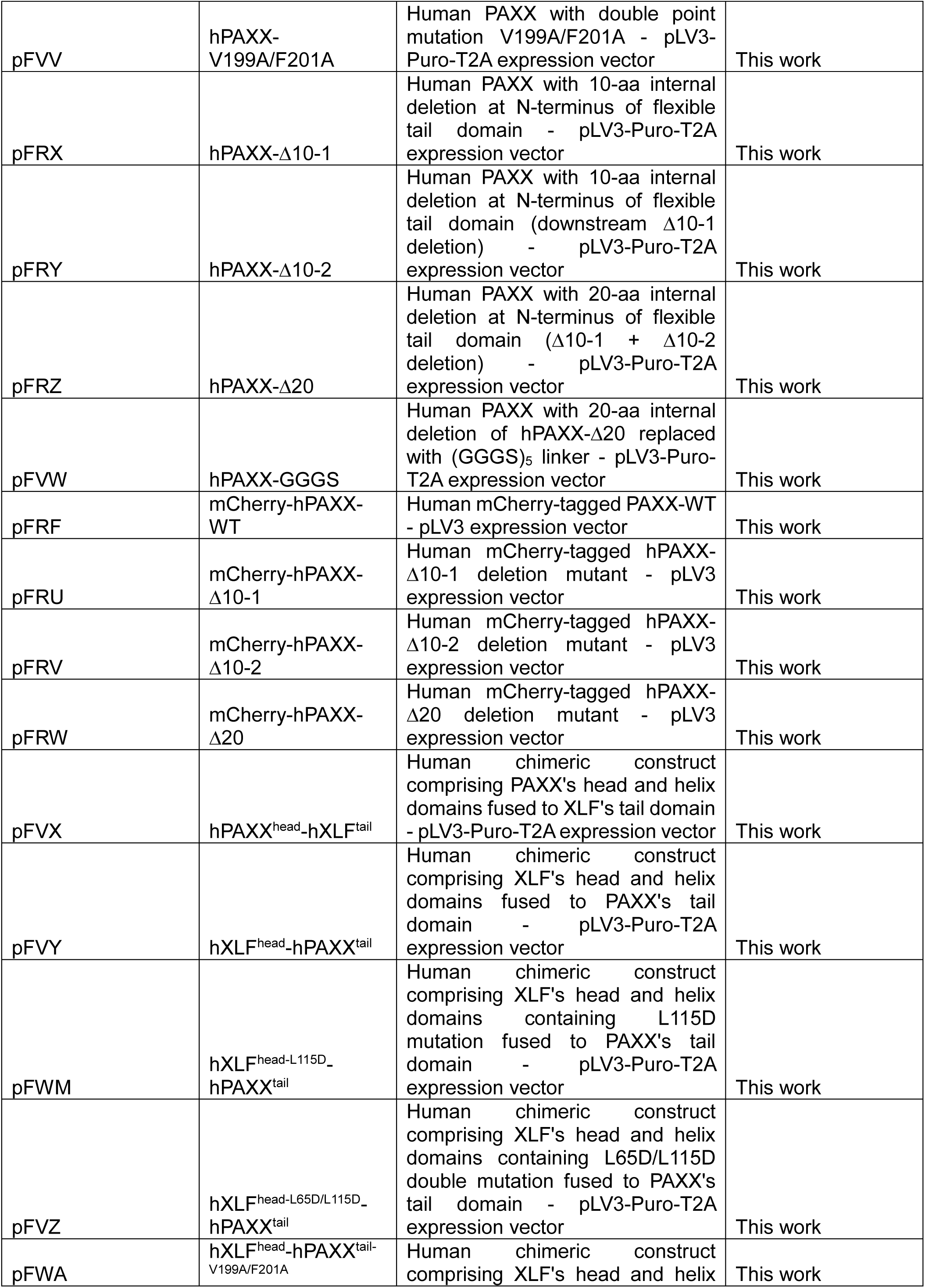

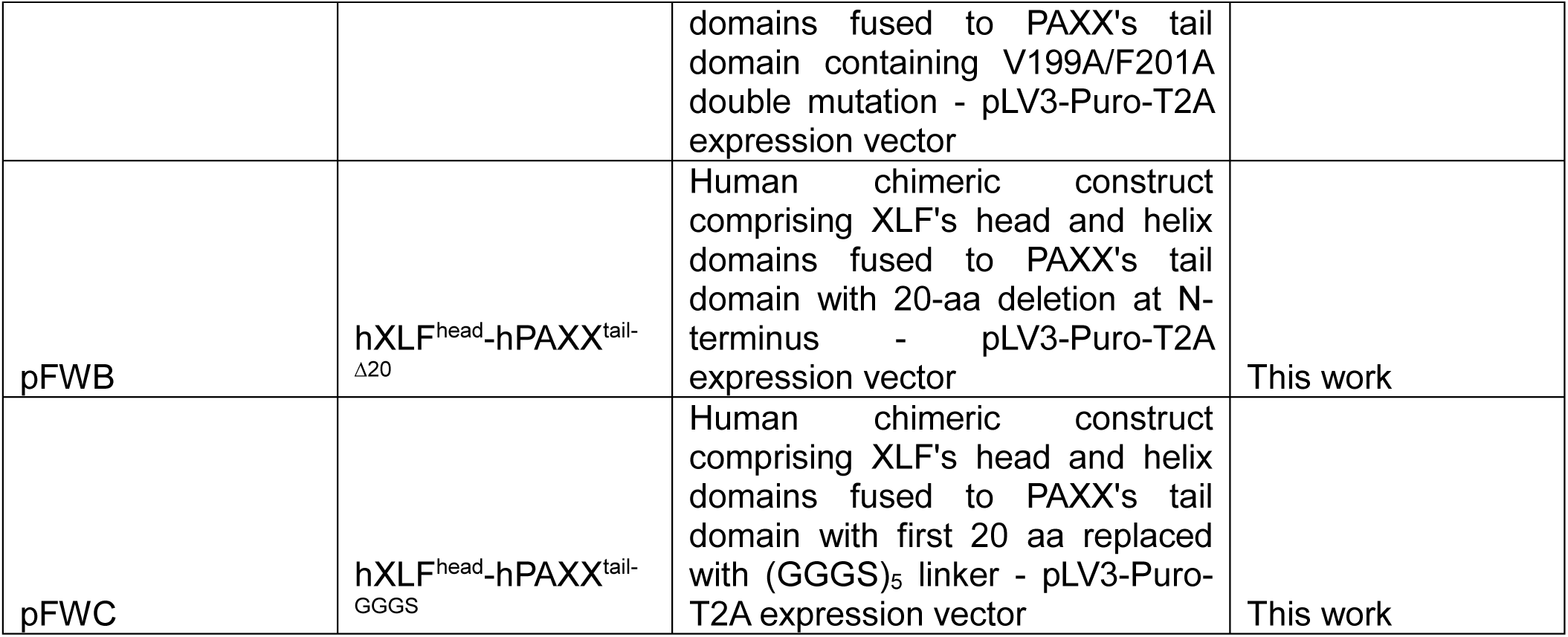

*Plasmids and DNA manipulations –* Xenopus *constructs.* All constructs were checked either by Sanger (Azenta) or nanopore (Plasmidsaurus) sequencing.

- pTG311: *X. laevis* cDNA was generated by reverse transcribing from *X. laevis* egg extract mRNA. An insert containing the sequence for *X. laevis* PAXX was PCR amplified from *X. laevis* cDNA using primers oTG574 and oTG575. This insert was inserted into a His-SUMO expression vector using NEBuilder HiFi assembly (New England Biolabs).
- pJZ001: pTG311 was PCR amplified using oligonucleotides oJZ001 and oJZ002 and then circularized using NEBuilder HiFi assembly (New England Biolabs).
- pJZ003: The backbone sequence was PCR amplified from pAM227 using oJZ003 and oJZ007. The insert was PCR amplified from pTG311 using oJZ004 and oJZ008. The two fragments were combined using NEBuilder HiFi assembly (New England Biolabs).
- pJZ018: The backbone sequence was PCR amplified from pTG311 using oJZ059 and oJZ030. The insert was PCR amplified from pTG311 using oJZ031 and oJZ058. The two fragments were combined using NEBuilder HiFi assembly (New England Biolabs).
- pJZ027: pTG311 was PCR amplified using oligonucleotides oJZ120 and oJZ121 and then circularized using NEBuilder HiFi assembly (New England Biolabs).
- pJZ028: pTG311 was PCR amplified using oligonucleotides oJZ122 and oJZ123 and then circularized using NEBuilder HiFi assembly (New England Biolabs).
- pJZ032: pTG311 was PCR amplified using oligonucleotides oJZ154 and oJZ155. The Sequence Manipulation Suite^50^ was used to generate a shuffled sequence of the first 20 residues of PAXX’s tails and the gene block gbJZ155 containing this sequence was synthesized. The backbone and gene block were combined using NEBuilder HiFi assembly (New England Biolabs).
- pJZ033: pTG311 was PCR amplified using oligonucleotides oJZ154 and oJZ155. The Sequence Manipulation Suite^50^ was used to generate another shuffled sequence distinct from the shuffled sequence in pJZ032 of the first 20 residues of PAXX’s tails and the gene block gbJZ156 containing this sequence was synthesized. The backbone and gene block were combined using NEBuilder HiFi assembly (New England Biolabs).
- pJZ034: The backbone sequence was amplified from pTG311 using oligonucleotides oJZ154 and oJZ155. The backbone and gbJZ157 were combined using NEBuilder HiFi assembly (New England Biolabs).
- pJZ035: pJZ027 was PCR amplified using oJZ150 and oJZ151 and then circularized using NEBuilder HiFi assembly (New England Biolabs).
- pJZ040: The backbone sequence was PCR amplified from pTG311 using oJZ059 and oJZ002. The insert was PCR amplified from pJZ001 using oJZ001 and oJZ058. The two fragments were combined using NEBuilder HiFi assembly (New England Biolabs).
- pJZ041: The backbone sequence was PCR amplified from pJZ001 using oJZ168 and oJZ030. The insert was PCR amplified from pTG311 using oJZ031 and oJZ058. The two fragments were combined using NEBuilder HiFi assembly (New England Biolabs).
- pJZ042: The backbone sequence was PCR amplified from pTG311 using oJZ173 and oJZ174. The insert was PCR amplified from pTG296 using oJZ175 and oJZ176. The two fragments were combined using NEBuilder HiFi assembly (New England Biolabs).
- pJZ043: The backbone sequence was PCR amplified from pTG440 using oJZ169 and oJZ170. The insert was PCR amplified from pTG311 using oJZ171 and oJZ172. The two fragments were combined using NEBuilder HiFi assembly (New England Biolabs).
- pJZ044: The backbone sequence was PCR amplified from pTG296 using oJZ169 and oJZ170. The insert was PCR amplified from pTG311 using oJZ171 and oJZ172. The two fragments were combined using NEBuilder HiFi assembly (New England Biolabs).
- pJZ055: pJZ027 was PCR amplified using oligonucleotides oJZ017 and oJZ018 and then circularized using NEBuilder HiFi assembly (New England Biolabs).
- pJZ059: pTG296 was PCR amplified using oligonucleotides oJZ017 and oJZ229 and then circularized using NEBuilder HiFi assembly (New England Biolabs).

### Plasmids and DNA manipulations – human cells

Expression vectors for untagged wild-type XLF and XLF-L115D mutant were obtained by PCR amplification of the corresponding human wild-type and mutant XLF cDNAs^34^ with XLF-Box-F and XLF-Mlu-R primers, followed by Hot-Fusion insertion^63^ between the BoxI and MluI restriction sites of pLV3-Puro-T2A^64^ i.e. downstream a puromycin resistance gene and a sequence encoding the T2A ribosomal skipping peptide. The expression vector for XLF-L65D-L115D double mutant was obtained in a similar manner with an additional step of PCR mutagenesis on pLV3-Puro-T2A-XLF-L115D as a template with XLF-L65D-F and XLF-L65D-R oligonucleotides as mutated inner primers (see below the list of primers) prior to two-fragment Hot-Fusion insertion. Wild-type XLF cDNA was PCR-amplified with XLF-Box-F and XLF-Mlu-R primers and inserted into Kpn2I and BcuI restriction sites of the pLV3-GFP plasmid^64^ to allow expression of GFP-tagged XLF.

Expression vectors for untagged wild-type PAXX and the PAXX-F201A and PAXX-V199A-F201A mutants were obtained by PCR amplification of PAXX cDNA^34^ with primers PAXX-Box-F and either PAXX-Bcu-R, PAXX-F201A-R or PAXX-VF>AA-R, respectively, followed by Hot-Fusion insertion between the BoxI and BcuI restriction sites of pLV3-Puro-T2A. The expression vectors for PAXX-Δ10-1, PAXX-Δ10-2, PAXX-Δ20 and PAXX-GGGS deletion mutants were obtained in a similar manner with an additional step of PCR mutagenesis with PAXX-Box-F and PAXX-Bcu-R as outer primers and the corresponding PAXX-mut-F and PAXX-mut-R oligonucleotides as mutated inner primers (see below the list of primers) prior to two-fragment Hot-Fusion insertion. Expression vectors for mCherry-tagged PAXX constructs were obtained as previously described^64^.

The vector encoding the PAXX^Head^-XLF^Tail^ chimera was assembled from separate PCR amplifications of PAXX^Head^ and XLF^Tail^ cDNAs with PAXX-Box-F/PAXX-xlf-R and paxx-XLF-F/XLF-Mlu-R pairs of primers, respectively, followed by a two-fragment Hot-Fusion insertion between the BoxI and MluI restriction sites of pLV3-Puro-T2A. Vectors encoding the various XLF^Head^-PAXX^Tail^ chimeras were assembled similarly from separate PCR amplifications of XLF^Head^ and PAXX^Tail^ cDNAs using suitable templates with XLF-Box-F/XLF-paxx-R and xlf-PAXX-F/PAXX-Bcu-R pairs of primers, respectively, with the following exceptions: xlf-PAXX-F/PAXX-VF>AA-Bcu-R pair of primers for the PAXX^Tail^-VF>AA fragment; XLF-Box-F/XLF-paxx-d20-R and xlf-PAXX-Δ20-F/PAXX-Bcu-R pairs of primers for the XLF^Head^-WT and PAXX^Tail^-Δ20 overlapping fragments, respectively; XLF-Box-F/XLF-paxx-Δ20+-R and xlf-PAXX-Δ20+-F/PAXX-Bcu-R pairs of primers for the XLF^Head^-WT and PAXX^Tail^-GGGS overlapping fragments, respectively.

All oligonucleotides were purchased from Eurofins Genomics (Ebersberg, Germany). Restriction and modifying enzymes (Phusion and T4 DNA Ligase) were from ThermoFisher Scientific (Illkirch, France). All constructs were checked by sequencing (Eurofins Genomics).

*Purification of* X. laevis *protein constructs*

All constructs except for XRCC4/LigIV^CD^ and all fluorescently labeled ybbR tagged constructs were purified according to the following protocol. These constructs were all H10-SUMO tagged. Expression plasmids were transformed into *E. coli* BL21(DE3)pLysS competent cells and plated on LB-Agar plates. Colonies were selected and grown overnight at 37°C with shaking in 5 mL LB cultures, which were used to inoculate large scale 1 L LB cultures. All cultures and plates contained Ampicillin at a concentration of 100 mg/L. Large scale cultures were grown at 37°C with shaking to an OD_600_ of approximately 0.6-0.8, then induced with IPTG to a final concentration of 1 mM. Induced cultures were incubated overnight at 16°C with shaking, harvested by centrifugation at 5000 RCF for 15 minutes, flash frozen in liquid nitrogen, and stored at - 80°C until commencement of subsequent purification steps. Pellets were thawed, weighed, and resuspended in 5 mL of lysis buffer (20 mM Tris pH 8, 400 mM NaCl, 10 mM Imidazole, 10% Glycerol v/v, 5 mM β-mercaptoethanol, 0.2 mM PMSF, and 1 Roche cOmplete Protease Inhibitor Cocktail tablet per 200 mL buffer) per gram of pellet. Resuspended pellets were Dounce homogenized, lysed using a Microfluidics LM10 Microfluidizer, and centrifuged at 47850 RCF at 4°C for 80 minutes. The lysate was incubated with rotation at 4°C with 1 mL Ni-NTA agarose beads (Qiagen) that had been equilibrated with lysis buffer. The beads were placed in a column, washed with 3 × 10 mL lysis buffer, then eluted with 7 × 1 mL elution buffer (20 mM Tris pH 8, 400 mM NaCl, 250 mM Imidazole, 10% Glycerol v/v, 5 mM β-mercaptoethanol). H6-Ulp1 protease was added to the eluate (final concentration of ∼10 µg/mL) to cleave off the H10-SUMO tag, and the eluate was dialyzed overnight in dialysis buffer (20 mM Tris pH 8, 400 mM NaCl, 10 mM Imidazole, 10% Glycerol v/v, 5 mM β-mercaptoethanol). The next day, the dialyzed protein was incubated with rotation at 4°C with 1 mL Ni-NTA agarose beads that had been equilibrated with dialysis buffer. After this incubation, the flowthrough was collected. This step bound the cleaved H10-SUMO tag and H6-Ulp1 protease to the beads while retaining the cleaved protein construct of interest in solution. The flowthrough was concentrated to 0.5 mL in volume, diluted in 9.5 mL of SP dilution buffer (20 mM Mes, 20 mM NaCl, 10% w/v Sucrose, 1 mM DTT, and 0.5 mM EDTA), and loaded onto an SP Sepharose Fast Flow column (GE Healthcare) that had been equilibrated with 10 column volumes of SP binding buffer (20 mM Mes, 10 mM NaCl, 10% w/v Sucrose, 1 mM DTT, and 0.5 mM EDTA) for cation exchange. The column was washed with 10 column volumes of SP binding buffer, and protein was eluted using a gradient of SP elution buffer (20 mM Mes, 1 M NaCl, 10% w/v Sucrose, 1 mM DTT, and 0.5 mM EDTA) and SP bind buffer. The elution gradient proceeded in two linear segments, with the first segment starting at 0% SP elution buffer (100% SP bind buffer) with target of 60% SP elution buffer over 20 column volumes and the second segment with target of 100% SP elution buffer over 5 column volumes. Peak fractions were pooled, loaded onto a Superdex® 200 Increase 10/300 GL size exclusion chromatography column (Cytiva), and eluted with size exclusion buffer (20 mM Tris pH 7.4, 300 mM sodium acetate pH 7.4, 1 mM DTT, 0.5 mM EDTA, and 10% Glycerol v/v) at a flow rate of 0.4 mL per minute. Peak fractions were pooled, concentrated to >10 μM as necessary using a 10000 MWCO spin concentrator (Amicon), aliquoted, flash frozen in liquid nitrogen, and stored at-80°C.

Fluorescently labeled ybbR tagged constructs were purified according to the above protocol up until the end of the cation exchange step. Peak fractions from cation exchange were pooled, buffer exchanged into labeling buffer (20 mM Tris pH 8, 300 mM NaCl, 10 mM MgCl2, 1 mM DTT, and 10% v/v Glycerol), and spin concentrated to >100 µM. Labeling reactions consisted of 50 µM ybbR-tagged protein in labeling buffer (100 µM ybbR tag concentration, as all constructs purified were homodimeric), 150 mM CoA-Cy5 dye, and 5 mM Sfp Synthase. The reaction volume was adjusted to 100 µL using labeling buffer. The labeling reaction was incubated overnight with rotation at 4°C. Size exclusion chromatography was performed on the labeling reaction using the protocol described above to futher purify the fluorescently labeled protein of interest and to separate out free dye. Peak fractions were pooled, aliquoted, concentrated, flash frozen in liquid nitrogen, and stored at-80°C.

For XRCC4/Lig4^K278R^, protein expression steps in *E. coli* and cell harvesting were identical to expression steps for all other constructs, except that large scale cultures were grown in TB (IBI scientific) prepared according to the manufacturer’s instructions and induced at an OD_600_ of ∼1.2. This construct bore both an N-terminal H6 tag and a C-terminal TwinStrep tag, allowing for dual affinity purification. Pellets were resuspended in 20 mL of XRCC4/Lig4 lysis buffer (20 mM Tris pH 8, 400 mM NaCl, 10 mM Imidazole, 10% Glycerol v/v, 1 mM DTT) per liter of cells, and 1 Roche cOmplete Protease Inhibitor Cocktail tablet per 2-3 liters of cell culture was added. Cells were lysed by sonication and centrifuged for 1 hour at 20000 RCF. The lysate was incubated with rotation at 4°C with 2 mL Ni-NTA agarose beads (Qiagen) that had been equilibrated with XRCC4/Lig4 lysis buffer. The beads were placed in a column, washed with 2 × 10 mL XRCC4/Lig4 lysis buffer, then protein was eluted with 10 × 1 mL XRCC4 elution buffer (20 mM Tris pH 8, 400 mM NaCl, 250 mM Imidazole, 10% Glycerol v/v, and 1 mM DTT). 2 mL bed volume of StrepTactin XT resin (IBA Lifesciences) was equilibrated in a column with XRCC4/Lig4 lysis buffer, the Ni-NTA eluate was applied to the StrepTactin column by gravity flow, the StrepTactin resin was washed with 2 × 10 mL XRCC4/Lig4 lysis buffer, and protein was eluted with 10 x 1 mL 1x buffer BXT (IBA Lifesciences) supplemented with 1 mM DTT. The eluate was dialyzed overnight into XRCC4/Lig4 SEC buffer (20 mM HEPES pH 7.5, 150 mM NaCl, 10% Glycerol v/v, and 1 mM DTT). Size exclusion chromatography was performed as described above using the XRCC4/Lig4 SEC buffer. Peak fractions were pooled, aliquoted, spin concentrated to 10 μM as necessary, flash frozen in liquid nitrogen, and stored at-80°C.

### Western blotting, antibodies and immunodepletion – egg extract

Anti-XLF and anti-XRCC4 antibodies were prepared as described previously^8^. Anti-PAXX antibody was raised by immunogenizing rabbits to the full length PAXX protein (Pocono farms). Generation of an antigen column and affinity purification of PAXX antibody were performed as described previously^8^.

To perform the protein depletions indicated in the main text, prepared antibodies were incubated with rProtein A Sepharose FastFlow (Cytiva) beads at a ratio of 4 µg antibody per µL of beads for anti-PAXX and 3 µg antibody per µL of beads for anti-XLF and anti-XRCC4; incubation occurred for at least 1 hour and up to overnight at 4°C with rotation. Bead-antibody slurries were then spun at 2000 RCF for ∼10 seconds to pellet beads, and the supernatant was aspirated off with a fine-tipped pipette tip. Beads were then washed, spun, and aspirated four times with 1x ELB (10 mM HEPES (pH 7.7), 50 mM KCl, 2.5 mM MgCl_2_) supplemented with 0.02% (v/v) Tween-20 and 94 g/L sucrose; the volume of wash buffer added during each wash was 10 times the volume of beads.

Extracts were supplemented with Nocodazole to a final concentration of 7.5 ng/µL prior to depletion and centrifuged for 10 minutes at 4°C at a speed of ∼17000 RCF (this was also done for experiments in which no depletion of the extract occurred). After centrifugation, any pellet that was present at the bottom of the tube of extract was removed. For each round of depletion, extract was added to antibody beads at a ratio of five parts extract to one part beads by volume and incubated with rotation at 4°C for 1 hour. At the end of each round of depletion, extract was aspirated off the beads, retained, and either added to additional beads at the above ratio for an additional round of depletion or to an empty tube for storage; extracts were either used immediately for assays or flash frozen in liquid nitrogen and stored at-80°C until later use. Mock depletions were performed as described previously^8^. Two rounds of depletion were performed for all depletions.

Extract samples for western blotting were treated with Lambda protein phosphatase to separate out the PAXX band from an overlapping nonspecific band. Lambda protein phosphatase treatment was carried out by incubating 2 volumes of egg extract, 1 volume of 10 mM MnCl_2_ (New England Biolabs), 1 volume of 10x NEBuffer for Protein MetalloPhosphatases (New England Biolabs), 1 volume of 400000 U/ml Lambda Protein Phosphatase (New England Biolabs), and 5 volumes of water at 30°C for 30 minutes. Phosphatase-treated samples were diluted 1:1 with 2x Laemmli buffer (Bio-Rad) and heated for 1 minute at 95°C. 5 µL of each sample was loaded per lane of a 15-well 4%-15% gradient polyacrylamide gel (Bio-Rad) and ran at 200 V for 50 minutes. Gels were transferred onto PVDF membranes (1 hour at 100 V and 4°C), which were blocked with 5% non-fat dry milk powder in PBST (w/v) for 30 minutes at room temperature and rinsed three times with PBST. Primary antibodies were added to 2.5% BSA and 0.04% Sodium Azide in PBST (1:1000 dilution for both antibodies), and membranes were incubated with primary antibody dilutions overnight at 4°C with shaking. Primary antibody was washed away with 3 x 5-minute rinses in PBST, after which the HRP-conjugated goat-anti-rabbit secondary antibody (Jackson ImmunoResearch) diluted 1:10000 in 5% non-fat dry milk powder in PBST was applied and incubated with the membrane at room temperature for 1 hour. Secondary antibody was washed away with 3 x 5-minute rinses in PBST, WesternBright ECL spray (Advansta) was applied and allowed to incubate with the membrane for ∼1 minute, and the membrane was imaged by chemiluminescence.

### End joining substrate generation for ensemble NHEJ assays

pJZ022 was linearized by digestion with NotI and EcoRI restriction enzymes (New England Biosciences) and gel purified. oJZ088 and oJZ109 were radiolabeled with ^32^P using T4 PNK (New England Biosciences). Labeling reactions consisted of 0.5 µM oligonucleotide, 0.5 µM [γ-32P] dATP (Perkin-Elmer), 2 U/µl T4 PNK (New England Biosciences), and 1X T4 PNK buffer (New England Biosciences). Reactions were incubated at 37°C for 30 minutes. To ensure complete phosphorylation of oligonucleotides, 50 µM cold dATP was added to phosphorylation reactions at the end of the 30 minute incubation, and the reaction was incubated for an additional 15 minutes at 37°C. oJZ033, oJZ088 and oJZ109 were cold phosphorylated using T4 PNK. Phosphorylation reactions consisted of 10 µM oligonucleotide, 400 µM dATP, 2 U/µl T4 PNK, and 1X T4 PNK buffer. Reactions were incubated at 37°C for 30 minutes. All phosphorylation reactions were stopped by the addition of EDTA to 25 mM and incubating at 65°C for 20 minutes. Oligonucleotides were annealed to generate forked and blunt ended adaptors for the substrate: hot and cold phosphorylated oJZ088 were annealed to oJZ089 to generate the forked-ended adaptor, hot and cold phosphorylated oJZ109 were annealed to oJZ110 to generate one set of blunt-ended adaptors, and cold phosphorylated oJZ033 was annealed to oJZ034 to generate another set of blunt-ended adaptors. The final concentrations of the annealed radiolabeled adaptors was 0.454 µM, and the final concentrations of the annealed cold phosphorylated adaptors was 8.7 µM. Adaptors were ligated to linearized pJZ022 backbone using T4 Ligase (New England Biosciences). The linearized backbone (25 nM), radiolabeled oJZ088/oJZ089 adaptor (for forked end substrate) or radiolabeled oJZ109/oJZ110 adaptor (for double blunt end substrate) (100 nM for both), cold phosphorylated oJZ088/oJZ089 adaptor (for forked end substrate) or cold phosphorylated oJZ109/oJZ110 adaptor (for double blunt end substrate) (300 nM for both), cold phosphorylated oJZ033/oJZ034 adaptor (400 nM), T4 Ligase (4 U/µl), and 1X T4 Ligase buffer (New England Biosciences) were incubated at 16°C for at least 1 hour and up to overnight and gel purified. Both hot-and cold-phosphorylated oJZ088/oJZ089 and oJZ109/oJZ110 duplexes were employed to ensure complete ligation of adaptors to DNA ends while minimizing use of radioactive isotopes.

### Ensemble NHEJ assays

Depleted extracts, which made up 74% of the reaction mixture by volume, were supplemented with a 33x ATP regenerating system (65 mM ATP, 650 mM phosphocreatine, 160 ng/µL creatine phosphokinase) to 1x final concentration. A closed-circular “carrier DNA” (1 µg/µL stock) that is required for efficient end joining^8^ was added to the reactions to a concentration of 30 ng/µL. Recombinant proteins were added where indicated. Recombinant WT XLF and the chimeric PAXX-XLF fusion constructs were always added to a concentration of 50 nM, while PAXX constructs were always added to a concentration of 50 nM in the time course assays (as in Figures 1 and 5) and to the indicated concentrations in the titration assays (as in Figures 3 and 4). All reactions were supplemented with protein storage buffer as necessary (size exclusion buffer, as above) so that the combined volumes of added protein and protein storage buffer was constant across all conditions and was equal to 10% of the reaction mixture by volume. Reactions were initiated by the addition of radiolabeled DNA substrate (description and synthesis of substrates as above and in the main text) to a concentration of 2 nM at a volume that was 10% of the final reaction volume. Reactions were carried out at room temperature and stopped by the addition of a 2 µL aliquot of the reaction mixture to 6 µL of stop solution (80 mM Tris, pH 8.0, 8 mM EDTA, 0.13% phosphoric acid, 10% Ficoll, 5% SDS, and 0.2% bromophenol blue) at the desired time points (0, 30, 60, and 120 minutes for the time course assays, and a single 120-minute time point for the titration assays). Samples were run on a Tris-borate-EDTA 0.8% Agarose gel, pressed, dried onto a HyBond-XL nylon membrane (GE healthcare) using a slab drier, and exposed to a phosphorscreen. Exposed phosphorscreens were scanned using a Typhoon FLA 7000 imager (GE Healthcare) to visualize the radiolabeled DNA. Band intensities were quantified in ImageJ. All end joining assays except for the assays in Extended Data 1 and 2 were performed in triplicate.

### Single molecule imaging – substrate preparation

For measurements involving intermolecular synapsis and protein recruitment to single DNA ends, the surface-bound, donor labeled dsDNA (Cy3-dsDNA) was generated as described previously^8^. Additionally, for measurements of intermolecular synapsis, the acceptor labeled dsDNA (Cy5-dsDNA) in solution was generated by annealing the Cy5-labeled oligonucleotide oAM138 with the digoxygenin-labeled oligonucleotide oAM368.

For measurements involving intramolecular synapsis and protein recruitment to the synaptic complex, a 1000 bp substrate dsDNA was generated by PCR-amplifying the plasmid pAM166 using Cy3b-labeled oligonucleotide oAM472 and either A647-labeled (for measuring SRSC stability) or unlabeled (for measuring recruitment of fluorescently labeled proteins to the synaptic complex) oligonucleotide oAM473 as primers. To incorporate an internal biotin, this PCR product was nicked using Nt.BbvCI (New England Biosciences) to generate two single-strand breaks and then heated and subsequently cooled in the presence of an excess of the biotinylated oligonucleotide oAM469 to denature the segment of DNA between the cuts made by Nt.BbvCI and anneal oAM469. The substrate was then ligated using T4 Ligase and gel purified by electroelution.

### Single molecule imaging – microscope construction and flow cell channel preparation

Microscope and flow cell construction were as described previously^8^. Prior to imaging, a series of ∼100 reference images was taken by capturing 50 ms exposures under both donor and acceptor excitation of a slide mounted with a coverslip to which 0.1 µM TetraSpeck^TM^ microspheres (Invitrogen) had been adhered, shifting the position of the slide between each image. Each flow cell channel was briefly incubated with 0.2 mg/ml Streptavidin, which was then washed out with PBS. The donor-labeled surface bound substrate DNA was diluted in PBS and introduced into the flow cell. The exact concentration of substrate DNA used varied and could be adjusted based on the DNA spot density observed in the donor channel after donor-labeled DNA application. Excess DNA was washed out with egg lysis buffer (ELB; 10 mM HEPES, pH 7.7, 50 mM KCl, 2.5 mM MgCl_2_) followed by ELB containing an oxygen scavenging system (OSS) with triplet state suppressors (OSS consisting of final concentrations of 4 mM PCA and 0.08 µM PCD; triplet state suppressors and their final concentrations were 0.8 mM Trolox and 0.5 mM each of methyl viologen and ascorbic acid).

### Single molecule imaging – reaction compositions and recording of movies

To initiate the reaction, a reaction mixture was flowed into the channel shortly after imaging commenced. This reaction mixture varied between experiments. For the intermolecular synaptic stability measurements in Figures 2 and 4, the mixture consisted of 19 µL PAXX/XLF double-depleted egg extract, the aforementioned OSS and triplet state suppressors at the previously listed concentrations, 50 nM recombinant XLF, 30 ng/µL “carrier DNA,” the indicated concentration of recombinant PAXX or chimeric construct, 120 nM anti-digoxygenin Fab (Roche), and 4 nM Cy5-dsDNA. The same reaction mixture was used for the experiment in Figure 6D, except instead of adding 50 nM recombinant XLF to all reactions, only the indicated constructs were added to the reactions to a concentration of 50 nM. For the synaptic transition efficiency assay in Figure 2, the reaction mixture consisted of 19 µL undepleted egg extract supplemented with nocodazole to a final concentration of 7.5 ng/µL, the aforementioned OSS and triplet state suppressors at the previously listed concentrations, 30 ng/µL “carrier DNA,” 400 nM XRCC4/Lig4^K278R^, the indicated concentration of recombinant PAXX, 120 nM anti-digoxygenin Fab, and 4 nM Cy5-dsDNA. For the PAXX-XLF reciprocal stabilization assay in Figure 3, the reaction mixture consisted of 19 µL PAXX/XLF double-depleted egg extract, the aforementioned OSS and triplet state suppressors at the previously listed concentrations, recombinant proteins as indicated (20 nM for Cy5-XLF and Cy5-PAXX and 50 nM for unlabled XLF and PAXX), and 30 ng/µL “carrier DNA.” For the measurements of Cy5-PAXX and Cy5-PAXX^Δ20^ on single-and double-ended substrates in Figure 4G, the reaction mixture consisted of 19 µL undepleted egg extract supplemented with nocodazole to a final concentration of 7.5 ng/µL, the aforementioned OSS and triplet state suppressors at the previously listed concentrations, recombinant proteins as indicated (20 nM for all), and 30 ng/µL “carrier DNA.” For the SRSC stability measurements in Figure 2, the reaction mixture consisted of 19 µL PAXX/XLF/XRCC4 triple-depleted egg extract, the aforementioned OSS and triplet state suppressors at the previously listed concentrations, 400 nM XRCC4/Lig4^K278R^, 50 nM recombinant XLF, recombinant PAXX as indicated (50 nM), and 30 ng/µL “carrier DNA.” Reaction volumes were adjusted to 25 µl using size exclusion buffer and ELB as needed, keeping salt concentrations consistent across experiments. Imaging conditions for all experiments except for the SRSC stability measurements consisted of one frame of donor excitation (532 nm) followed by four frames of acceptor excitation (642 nm) at a frame rate of 5 frames per second. Surface laser power density was 600 mW/cm^2^ for the 532 nm laser and 300 mW/cm^2^ for the 642 nm laser as measured on the table. For the SRSC stability measurements, one frame of donor excitation (exposure time of 0.5 s) was followed by one frame of acceptor excitation (exposure time of 0.5 s), with a 2 s pause between each frame of excitation. Surface laser power density was 500 mW/cm^2^ for the 532 nm laser and 250 mW/cm^2^ for the 642 nm laser as measured on the table.

### Single molecule imaging – data analysis

Alignment of donor and acceptor channels was performed using a previously described method^65^. Briefly, a custom MATLAB script was used to identify spots in the 532 and 641 nm channels of the reference images by fitting to a 2D Gaussian. Spots in the 532 nm channel that were greater than 6 pixels from any other spot and also less than 6 pixels from the corresponding spot in the 641 nm channel were selected for calculating the alignment. The alignment was calculated using a previously described method. Briefly, the reference image spots in the 532 nm channel (x_1_, y_1_) were mapped onto the spots in the 641 nm channel (x_2_, y_2_) to identify a partner spot using the transformation

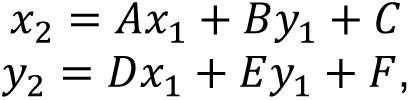

where A, B, C, D, E, and F are fit parameters. Different fit parameters are determined for each point (x_1_, y_1_) to account for the non-uniformity of the image. All points (x_1_, y_1_) that map to a point greater than 3 pixels from a partner point (x_2_, y_2_) are eliminated from the calibration list. The process is then repeated, decreasing the threshold for the distance between the 532 nm channel spots and their corresponding 641 nm channel partners to 2 pixels, then 1 pixel, then 0.5 pixels.

Custom MATLAB code was used to detect spots and integrate their intensities for each movie. Spots were first detected and integrated in the 532 nm channel; this was done by fitting spots to a 2D Gaussian. Each spot in the 532 nm channel was mapped to its corresponding location in the 641 nm channel, and the intensity of the spot in this location was also integrated. Stage drift was estimated by computing the average change in position between successive frames in the spots in the 532 nm channel, and this change in position was applied to all frames to correct for drift. Non-specific interactions with the coverslip glass were monitored by selecting points in the 532 nm channel that were at least 6 pixels away from any spots; these “dark spots” were also mapped to their corresponding location in the 641 nm channel. The intensities of each 532 nm spot or dark spot and their corresponding location in the 641 nm channel were integrated over time.

For all movies except the movies used in the experiments in Figures 2F and 2H-I, only the first 270 seconds of the movie were analyzed. For the movies used in the synaptic transition efficiency measurements in Figure 2F, the first 480 seconds were analyzed, and for the movies used in the SRSC stability measurements in Figure 2H-I, the first 870 seconds were analyzed. To detect colocalization of acceptor and donor signals, we used custom MATLAB code to analyze each trace generated by the above integration protocol. Each trace contained the donor and acceptor intensities under donor or acceptor excitation; these values were then normalized to a median value above a predetermined threshold. Colocalization events were defined by the acceptor signal meeting several criteria defined by a list of parameters including the minimum and maximum spot size, the maximum distance between the corresponding spots in the donor and acceptor channels, and the minimum normalized intensity and minimum change in the normalized signal for an increase in the acceptor signal to be considered a binding event. After a tentative set of parameters was selected, the parameters were adjusted by manually inspecting the traces and selecting parameters that minimized the number of detected nonspecific acceptor binding events while also maximizing the number of apparent acceptor binding events that were detected.

If necessary, the corrected FRET efficiency was calculated from the donor and acceptor signal intensities, as well as from information about donor fluorophore photobleaching (computed using MATLAB’s ischange() function). The intensity and signal change thresholds for detecting FRET events were adjusted based on manual inspection of the FRET traces. Only detected FRET events immediately following acceptor binding were considered for analysis. For measurements of LRSC stability, all acceptor binding events that transitioned to a high-FRET state were censored for the final Kaplan-Meier survival analysis. FRET was not calculated for any measurements involving the colocalization of acceptor-labeled protein to the synaptic complex, and no colocalization events were censored. For measurements of SRSC stability, all FRET events whose end coincided with a fluorophore photobleaching event (acceptor photobleaching also computed with the ischange() function for these measurements) or with the end of the analysis time frame were censored.

### Mass photometry

Mass photometry measurements were performed on a Refeyn TwoMP mass photometer according to manufacturer’s instructions. Briefly, a calibrant mixture containing β-Amylase and thyroglobulin was diluted to 1x concentration in PBS. The calibrant was imaged for 60 s, and peaks were assigned to the expected molecular weights of the species present to generate a calibration curve. This calibration curve was then used to calculate the molecular weights corresponding to the contrasts obtained from the measured constructs. All constructs measured using mass photometry were diluted to 15 nM and imaged for 60 s. Data was analyzed using Refeyn’s software.

### Molecular graphics

The molecular graphics in Extended Data 4A were generated using USCF ChimeraX^66^.

*Cell lines, cell culture and cell engineering.* Human osteosarcoma U2OS cells (ECACC, Salisbury, UK) and human embryonic HEK-293T cells (ATCC, Manassas, VA, USA), were grown in DMEM (Eurobio, France) supplemented with 10% fetal calf serum (Eurobio, France), 125 U/ml penicillin, and 125 μg/ml streptomycin. Cells were maintained at 37°C in a 5% CO_2_ humidified incubator.

U2OS cells knocked-out for *PAXX* and HEK-293T cells knocked-out for both *PAXX* and *NHEJ1* (*XLF*), were obtained using the CRISPR/Cas9 technology, as previously described^23^. The generation of modified U2OS and HEK-293T cell populations expressing various XLF and PAXX constructs was obtained as previously described^67^. Cell populations expressing untagged XLF or PAXX constructs were selected with 0.75 µg/ml puromycin for two weeks following transduction (InvivoGen, Toulouse, France).

### Western blotting and antibodies – human cells

Cell pellets were washed with phosphate-buffered saline (PBS) and resuspended in lysis buffer (50 mM HEPES-KOH, pH 7.5, 450 mM NaCl, 1 mM EDTA, 1% Triton X-100) supplemented with Halt protease inhibitor cocktail (ThermoFisher Scientific). Cells were lysed by four freeze/thaw cycles in liquid nitrogen and 37°C water bath. Lysates were cleared by centrifugation and protein concentrations were determined using the Bradford assay (Bio-Rad, Hercules, CA). Equal amounts of proteins were mixed with concentrated loading sample buffer to 1X final concentration (50 mM Tris.HCl pH 6.8, 10% glycerol, 1% SDS, 300 mM 2-mercaptoethanol, 0.01% bromophenol blue), heat-denatured, separated by SDS-PAGE on Miniprotean TGX stain-free 4-15% gradient gels (Bio-Rad, Hercules, CA) and blotted onto Immobilon-FL 0.45 µm PVDF transfer membranes (Millipore). Membranes were blocked for 60 min with 5% non-fat dry milk in PBS, 0.1% Tween-20 (Sigma-Aldrich) (PBS-T buffer), incubated as necessary with primary antibody diluted 1/1000 in PBS-T containing 1% bovine serum albumin (immunoglobulin-and lipid-free fraction V; Sigma-Aldrich) and washed 3 times with PBS-T. Membranes were incubated for 1 h with HRP-conjugated secondary antibodies (Jackson Immunoresearch Laboratories) diluted 1/10000 in PBS-T and washed five times with PBS-T. Immunoblots were visualized by enhanced chemiluminescence (Western Lightning Plus-ECL; Perkin Elmer) and autoradiography. Primary antibodies used: mouse monoclonal antibodies anti-Ku80 (clone 111; Thermo Fisher Scientific, MA5-12933, lot: WB31922872) and anti-Ku70 (clone N3H10; Thermo Fisher Scientific, MA5-13110, lot: WE27669780); rabbit polyclonal antibodies anti-XLF (Abclonal, A19957, lot: 00040701118) and anti-PAXX (Novus, NBP1-94172, lot: C118006).

### Multiphoton laser micro-irradiation

Live cell multiphoton laser micro-irradiation was conducted as previously described^34^.

### Cell imaging

To observe the subcellular localization of PAXX deletion mutants, U2OS cells expressing the different mCherry-tagged PAXX constructs were seeded in 35-mm glass-bottom culture dishes (MatTek) prior to imaging with an Olympus IX73 fluorescence microscope equipped with an Olympus UPlanFL N 40X/0.75 Ph2 objective lens. Images were taken with an Olympus DP26 camera.

### In vivo DNA end-joining assays

Direct end-joining activity was assessed as described previously^51^. In brief, HEK-293T cells were seeded to 20-40% confluence in 6-well plates and transfected 24 h later with a mix of Cas9-targeted NHEJ reporter substrate, Cas9/gRNA expressing vector and mCherry expressing plasmid as an internal control. Cells were trypsinized 2 days post-transfection, washed with PBS and analyzed by flow cytometry on a Fortessa X-20 cell analyzer (BD Biosciences). The integrated green fluorescence signal accounting for NHEJ-mediated repair events (% positive cells x mean fluorescence) was normalized to that of transfection efficiency (red fluorescence) (Extended Data 7).

### Declaration of generative AI and AI-assisted technologies in the writing process

During the preparation of this work, the authors used GPT-4 for editing and improving clarity of the text. After using this tool/service, the authors reviewed and edited the content as needed and take(s) full responsibility for the content of the published article.

## Data availability

Uncropped and unedited gel and blot images (for all experimental replicates, not just the representative gel images included in the figures), source data are available at https://doi.org/10.5281/zenodo.20292756.

## Code availability

Data and code for extracting relevant measurements from single molecule experiments are available at https://doi.org/10.5281/zenodo.20292756.

## Acknowledgements

We thank members of the Walter and Loparo laboratories for helpful discussion, Johannes Walter for discussion and access to his group’s frog facility, Marie Bao for feedback on the manuscript, Thomas Graham for cloning *X. laevis* PAXX, and the Center for Macromolecular Interactions at Harvard Medical School for supporting us in performing mass photometry experiments. Molecular graphics were generated using UCSF ChimeraX^66^, developed by the Resource for Biocomputing, Visualization, and Informatics at the University of California, San Francisco, with support from National Institutes of Health R01GM129325 and the Office of Cyber Infrastructure and Computational Biology, National Institute of Allergy and Infectious Diseases. This work was supported by grants from the National Institutes of Health R01GM115487 and R35GM158214 (to J.J.L) and T32GM008313 (to J.Z.). Additional support came from the French National Research Agency (ANR-20-CE11-0026 and ANR-24-CE44-1817) to N.B., P.C., P.F. and S.B.. P.F., N.B., P.C. and S.B. acknowledge the imaging facility TRI, member of the national infrastructure France-BioImaging infrastructure supported by the French National Research Agency (ANR-24-INBS-0005 FBI BIOGEN). P.C. is a scientist from INSERM.

## Author Contributions

J.Z. and J.J.L. conceived of the study and prepared the manuscript. J.Z., P.F., and N.B. performed experiments. J.J.L., S.B., and P.C. acquired funding. S.B. and P.C. provided ideas and contributed to the analysis of the data obtained from human cells. A.T.M. provided early versions of code for single molecule analysis, which J.Z. modified for the specific analyses performed in this work. All authors edited the manuscript.

