## Extended Data for "Sequential division of labor between PAXX and XLF drives NHEJ synaptic complex stability and maturation"

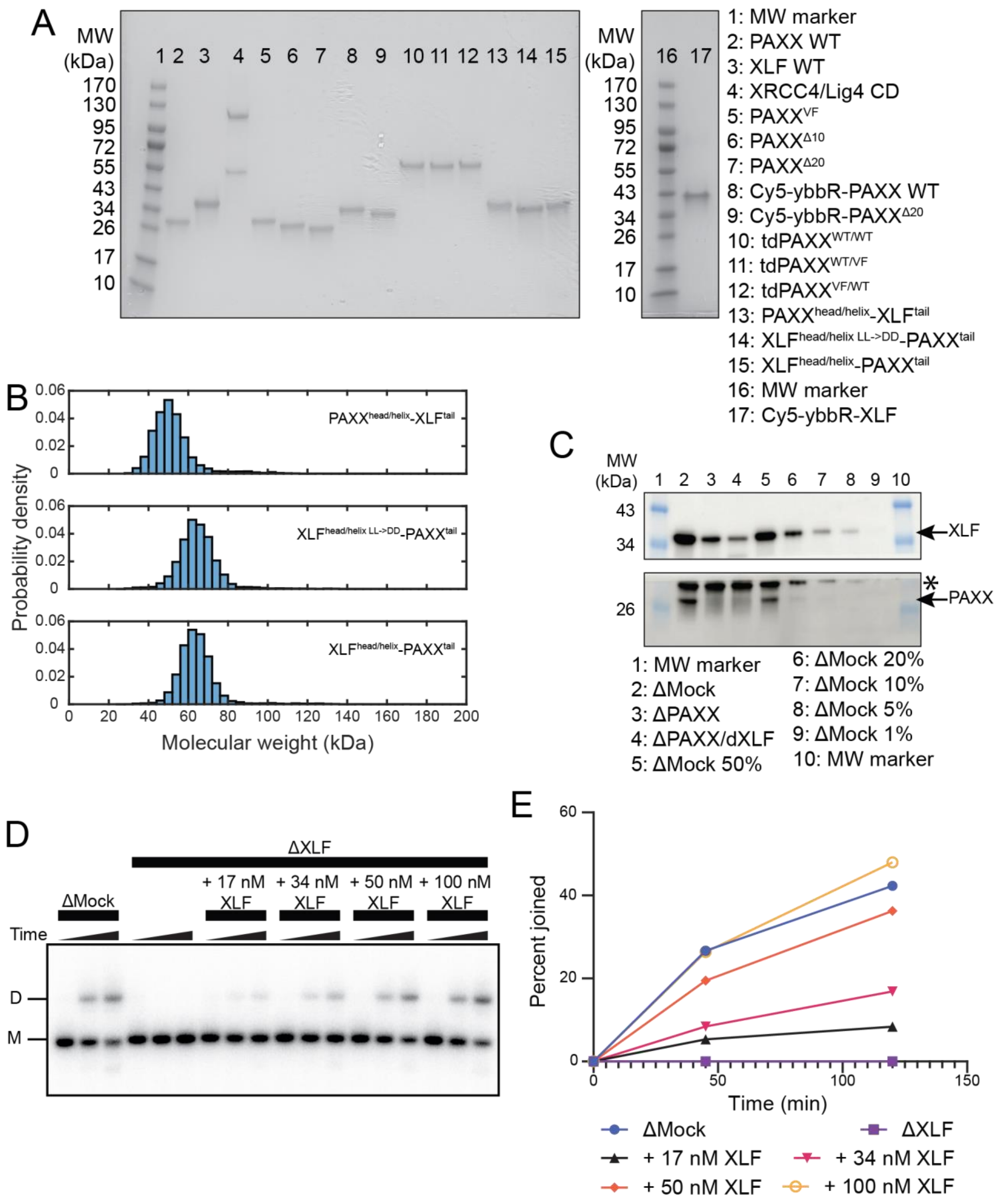


**Extended Data 1.** (A) Coomassie-stained SDS-PAGE gels of all purified *X. laevis* recombinant proteins used in this work. (B) The relative positions of the bands for the chimeric constructs in Extended Data 1A (lanes 13-15) are not consistent with their relative molecular weights (the $PAXX^{head/helix}$-$XLF^{tail}$ monomer is ~23 kDa, while the $XLF^{head/helix}$-$PAXX^{tail}$ and $XLF^{head/helix\ L\rightarrow D}$-$PAXX^{tail}$ monomers are both ~31 kDa). To verify the molecular weights and identities of the chimeric constructs, we performed mass photometry measurements to measure the molecular weights of these constructs. These measurements show peaks at ~50 kDa for $PAXX^{head/helix}$-$XLF^{tail}$ and ~60 kDa for $XLF^{head/helix}$-$PAXX^{tail}$ and $XLF^{head/helix\ L\rightarrow D}$-$PAXX^{tail}$, consistent with the expected molecular weights for each construct. (C) Western blots showing depletion of PAXX and XLF by anti-PAXX and anti-XLF antibodies, respectively, as well as partial depletion of XLF by anti-PAXX antibody. Top: anti-XLF primary. Bottom: anti-PAXX primary. Asterisk indicates location of nonspecific band. (D) End joining assay showing end joining time courses in XLF-depleted extract rescued with varying concentrations of recombinant XLF. End joining in depleted extract rescued with 50 nM XLF is comparable to end joining in mock-depleted extract, justifying our choice of this concentration for our XLF rescues. M: monomeric substrate, D: dimeric product. (E) Quantification of Extended Data 1D. Some information about depletion conditions has been omitted from the legend to save space.

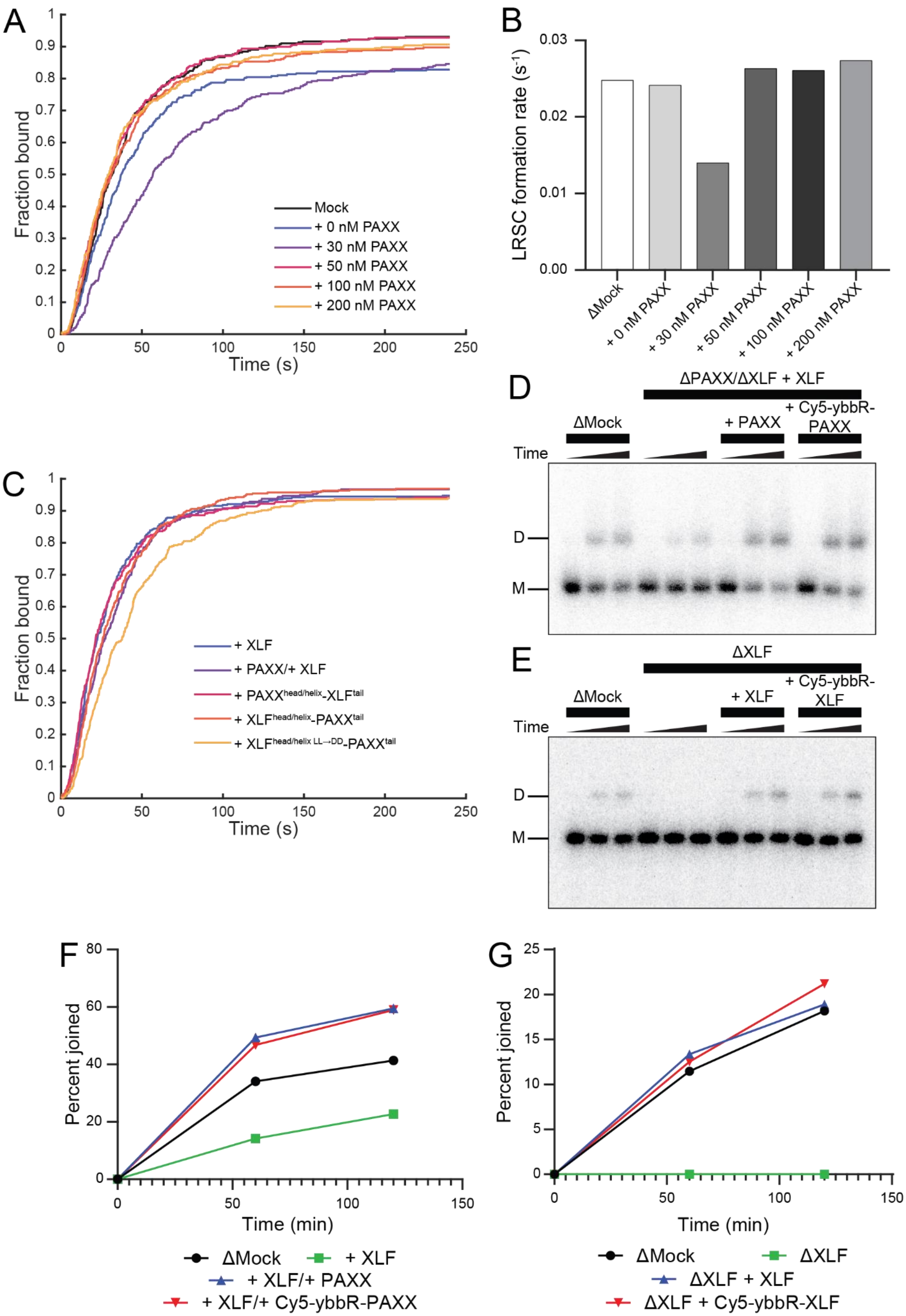


**Extended Data 2.** (A) Association of acceptor-labeled DNA to donor-labeled DNA over time for experiment described in Figure 2A-D. Association was measured by determining the fraction of donor-labeled, surface-bound dsDNAs that had experienced at least one donor-acceptor colocalization event at each timepoint. (B) Rate constants extracted from Extended Data 2A. (C) Association of acceptor-labeled DNA to donor-labeled

DNA over time for experiment described in Figure 6D. (D) End joining assay in egg extract with indicated depletions and rescues demonstrating that Cy5-ybbR-PAXX stimulates NHEJ to a comparable level as WT PAXX. (E) End joining assay in egg extract with indicated depletions and rescues demonstrating that Cy5-ybbR-XLF stimulates NHEJ to a comparable level as WT XLF. For Extended Data 2D-E, M indicates monomeric substrate, and D indicates dimeric product. (F) Quantification of Extended Data 2D. Some information about depletion conditions has been omitted from the legend to save space. (G) Quantification of Extended Data 2E.

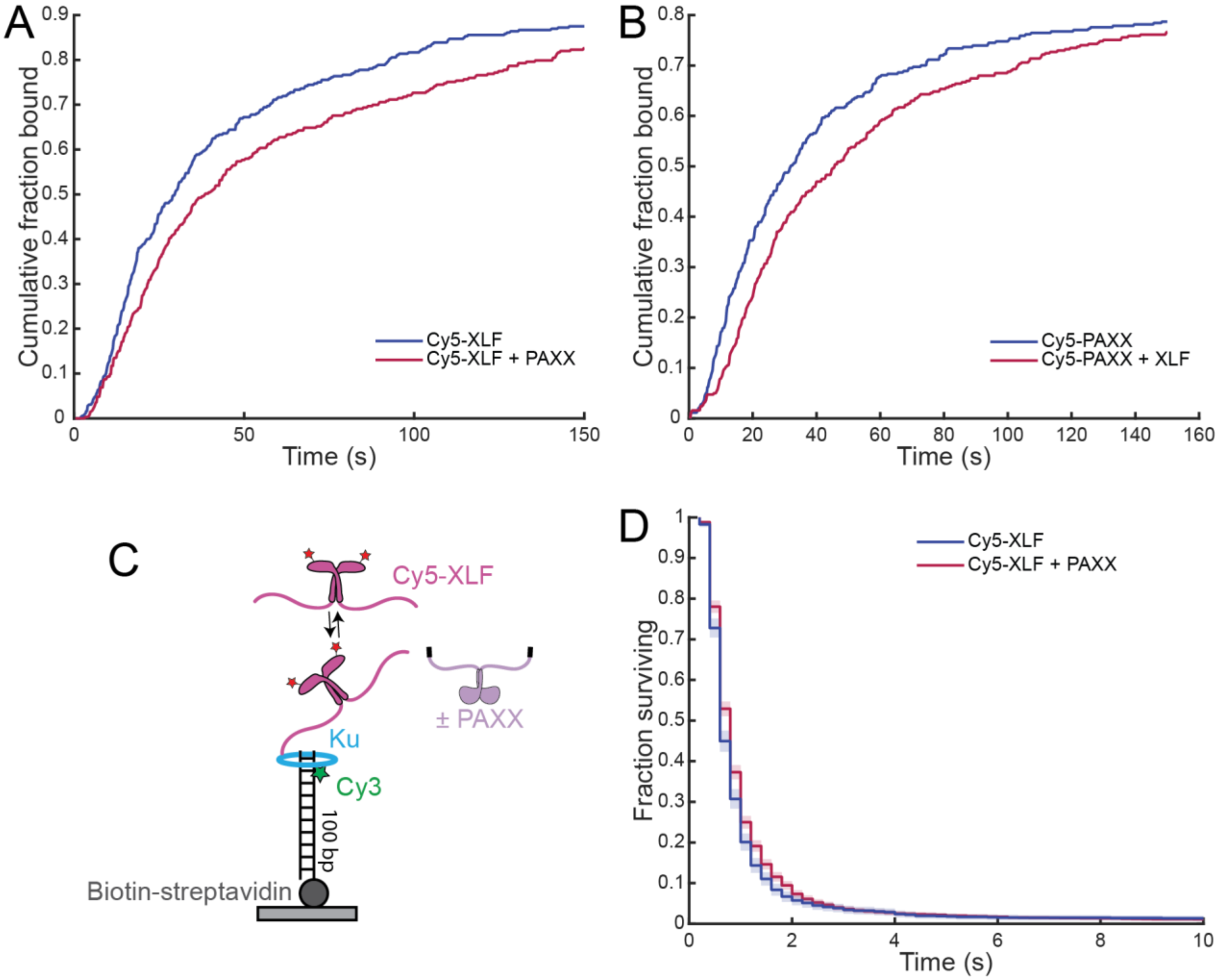


**Extended Data 3.** (A) Association of Cy5-XLF to DNA ends for experiment in Figure 3A. (B) Association of Cy5-PAXX to DNA ends for experiment in Figure 3B. (C) Substrate and proteins used to measure XLF stability on a single DNA end. (D) Kaplan-Meier survival plot showing stability of Cy5-XLF on single DNA end in the presence and absence of PAXX. Measurements were performed in PAXX/XLF double-depleted extract supplemented with the indicated proteins. Shaded regions on Kaplan-Meier curves represent 95% confidence interval.

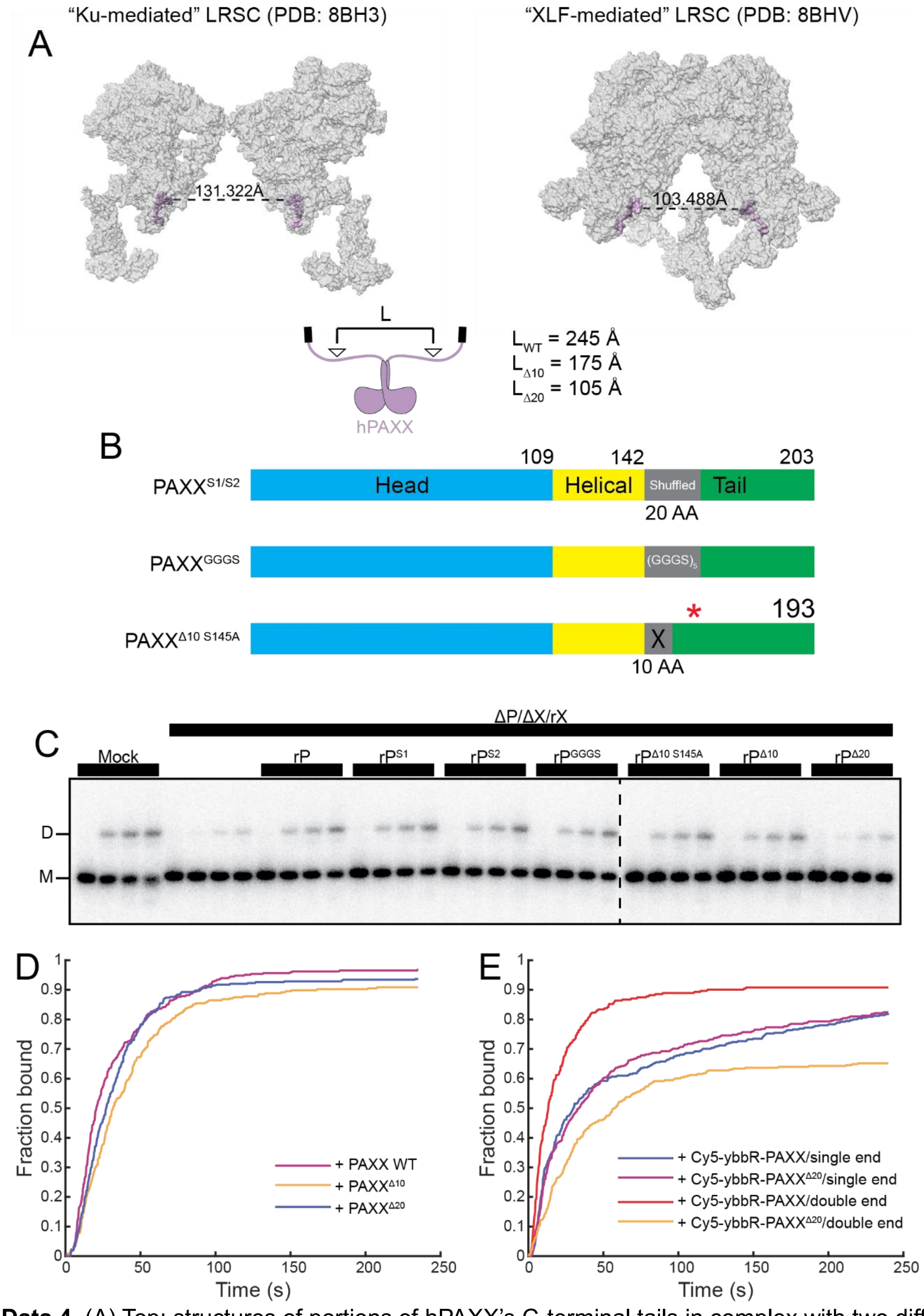


**Extended Data 4.** (A) Top: structures of portions of hPAXX's C-terminal tails in complex with two different LRSCs, along with measurements of the distances between the C180 residues present at the N-termini of the resolved portions of PAXX's tail domain. PAXX is colored in light purple and opaque; the rest of the structure is colored in grey and transparent. Bottom: schematic of distance estimation and estimates of the distances between the C180 (or the corresponding residue in the shortened tail constructs) in dimers of WT hPAXX, $hPAXX^{\Delta10}$, and $hPAXX^{\Delta20}$. Arrowheads indicate position of C180. Estimates were made by determining the number of amino acids between C180 (or corresponding residue) and the last N-terminal residue of the coiled-coil domain ($T145^{20}$) for each construct, multiplying by 2, and then multiplying by 3.5 Å per residue. (B) Domain architecture of scrambled and GGGS controls for Figure 4. (C) End joining in depleted extract rescued with XLF and PAXX constructs from (B). Results were run across two gels; dotted line indicates border between gels. M: monomeric substrate, D: dimeric product. (D, E) Association data from experiments in Figures 4E and 4G. Some information about depletion conditions has been omitted from the legend in Extended Data 4D to save space.

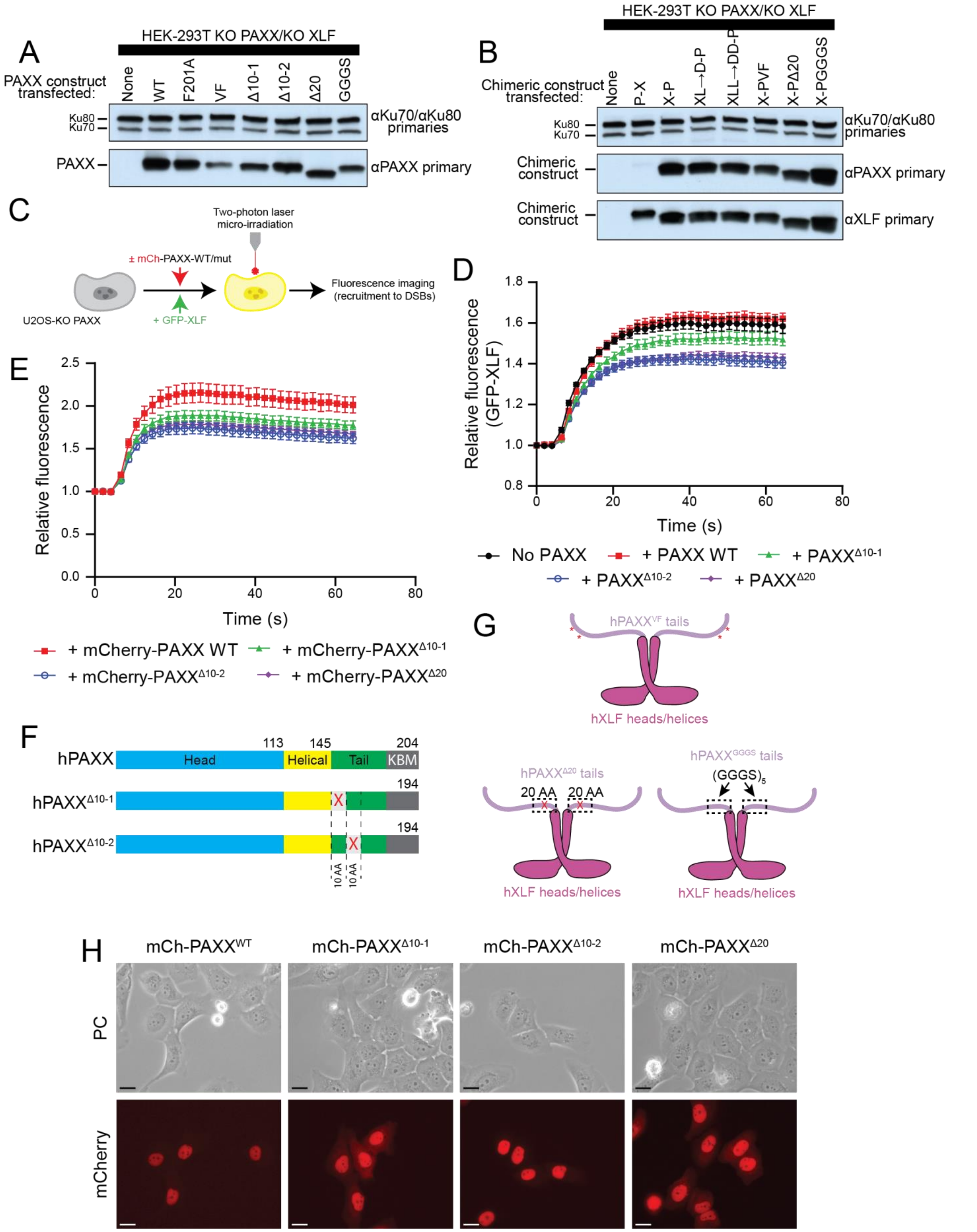


**Extended Data 5.** (A) Western blots showing expression of KBM mutant, shortened tail, and GGGS substitution PAXX constructs from Figure 4H in KO PAXX+XLF HEK-293T cells stably expressing the indicated construct. (B) Western blots showing expression of chimeric constructs from Figure 6F in KO PAXX+XLF HEK-293T cells stably expressing the indicated construct. (C) Schematic of laser irradiation experiment to measure recruitment of fluorescently labeled XLF and PAXX to sites of DNA damage. (D) Recruitment of GFP-XLF to laser irradiation sites in U2OS KO PAXX cells stably expressing GFP-XLF and the indicated PAXX construct. Error bars in Extended Data 5C-D represent ± SEM. (E) Recruitment of indicated mCherry-PAXX constructs in U2OS KO PAXX cells stably expressing the indicated PAXX construct. (F) Domain architecture of some human PAXX constructs used for the experiment in Figure 4H. (G) Cartoons of some human PAXX/XLF chimeric constructs used for the experiment in Figure 6F. (H) Nuclear localization of mCherry-labeled shortened tail PAXX constructs. Scale bars represent 20 μm.

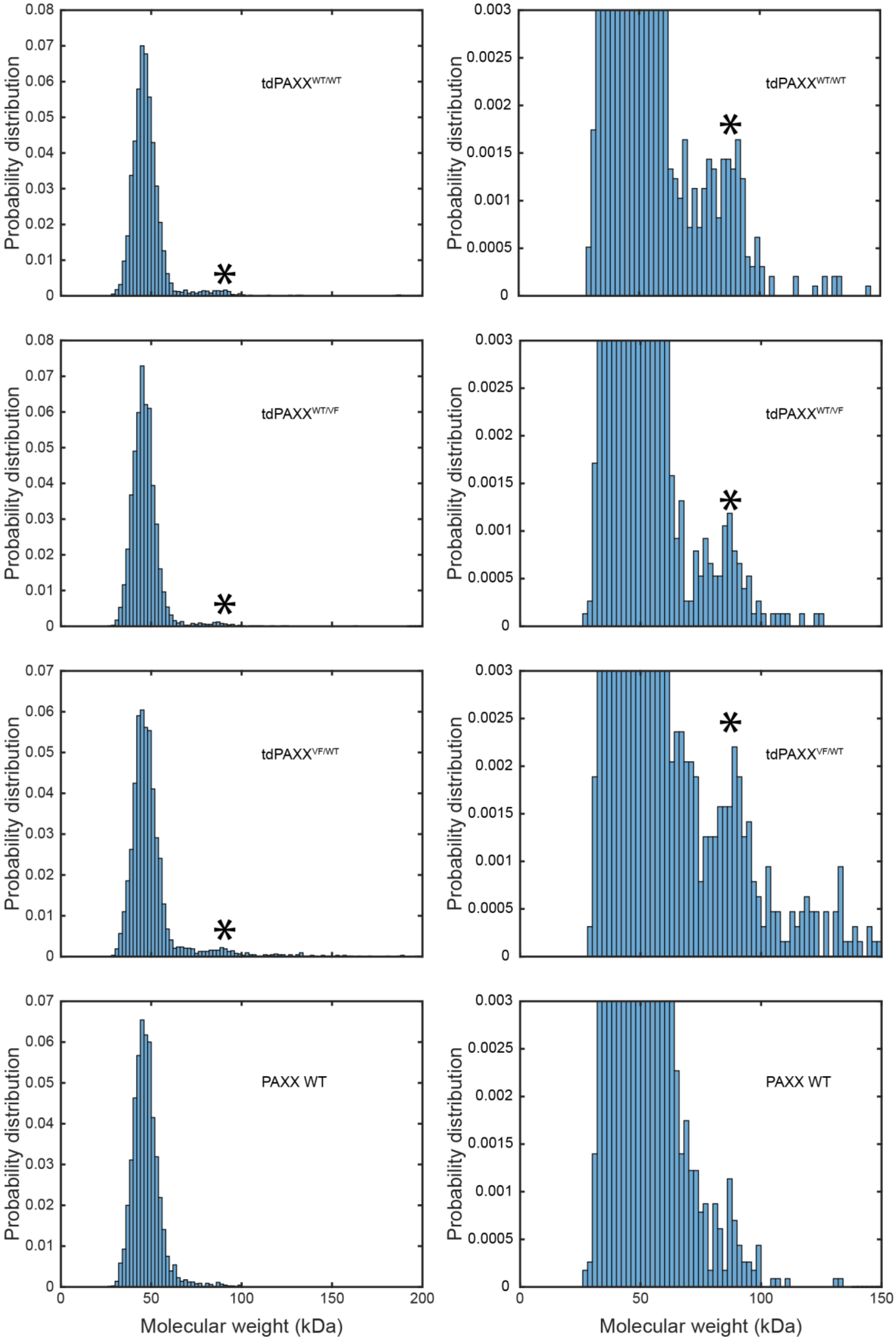


**Extended Data 6.** Mass photometry of tandem dimer constructs and WT PAXX. Asterisk indicates peak at ~80 kDa corresponding to the expected molecular weight of a dimer of tandem dimers. Histograms on right are zoomed-in plots of histograms on left.

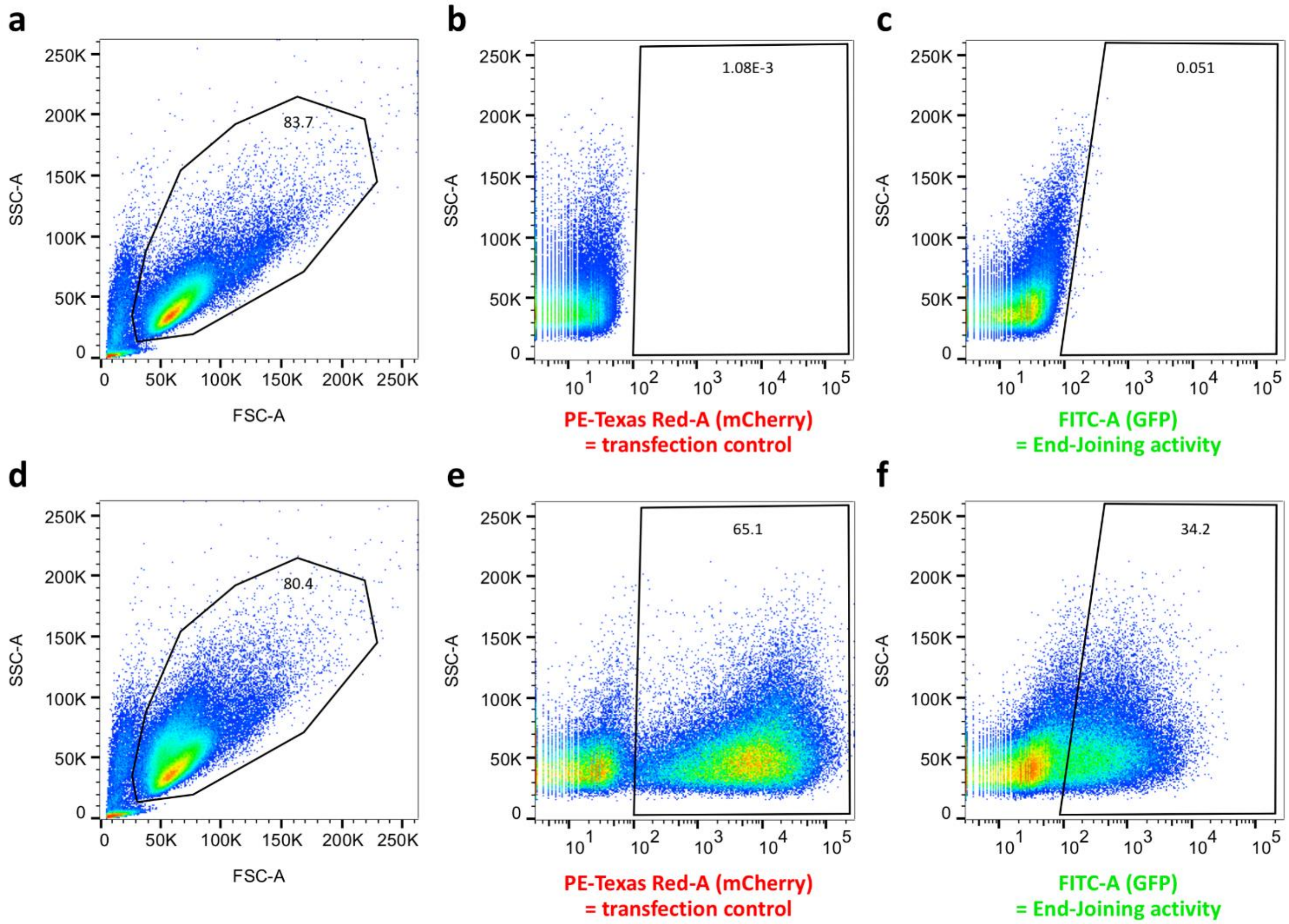


**Extended Data 7: flow cytometry gating strategy used for the cell-based end-joining assay.** (A) Cell debris and dead cells were excluded based on forward scatter area (FSC-A) and side scatter area (SSC-A). Gated FSC/SSC population of untransfected 293T cells (A) was used to position boundaries between negative and positive cells for red (PE-Texas Red-A) and green (FITC-A) fluorescence, (B) and (C), respectively. The same gates were applied to transfected cells (D, E, and F) to measure red and green fluorescence accounting for transfection efficiency and end-joining repair activity, respectively. Values correspond to the percentages of gated cells.
